# The quasi-universality of nestedness in the structure of quantitative plant-parasite interactions

**DOI:** 10.1101/2021.03.03.433745

**Authors:** Moury Benoît, Audergon Jean-Marc, Baudracco-Arnas Sylvie, Ben Krima Safa, Bertrand François, Boissot Nathalie, Buisson Mireille, Caffier Valérie, Cantet Mélissa, Chanéac Sylvia, Constant Carole, Delmotte François, Dogimont Catherine, Doumayrou Juliette, Fabre Frédéric, Fournet Sylvain, Grimault Valérie, Jaunet Thierry, Justafré Isabelle, Lefebvre Véronique, Losdat Denis, C. Marcel Thierry, Montarry Josselin, E. Morris Cindy, Omrani Mariem, Paineau Manon, Perrot Sophie, Pilet-Nayel Marie-Laure, Ruellan Youna

**Author notes:** Bayer Seeds SAS, Chemin de Roquemartine Mas Lamy, 13670, Saint-Andiol, France. Syngenta Seeds B.V., Westeinde 62, P.O. box 2, Enkhuizen 1600 AA, The Netherlands. **Cite as:** Moury B, Audergon J-M, Baudracco-Arnas S, Ben Krima S, Bertrand F, Boissot N, Buisson M, Caffier V, Cantet M, Chanéac S, Constant C, Delmotte F, Dogimont C, Doumayrou J, Fabre F, Fournet S, Grimault V, Jaunet T, Justafré I, Lefebvre V, Losdat D, Marcel TC, Montarry J, Morris CE, Omrani M, Paineau M, Perrot S, Pilet-Nayel M-L, Ruellan Y (2021) The quasi-universality of nestedness in the structure of quantitative plant-parasite interactions. bioRxiv, 2021.03.03.433745, ver. 4 recommended and peer-reviewed by PCI Evolutionary Biology. https://doi.org/10.1101/2021.03.03.433745.

## Abstract

Understanding the relationships between host range and pathogenicity for parasites, and between the efficiency and scope of immunity for hosts are essential to implement efficient disease control strategies. In the case of plant parasites, most studies have focused on describing qualitative interactions and a variety of genetic and evolutionary models has been proposed in this context. Although plant quantitative resistance benefits from advantages in terms of durability, we presently lack models that account for quantitative interactions between plants and their parasites and the evolution of these interactions. Nestedness and modularity are important features to unravel the overall structure of host-parasite interaction matrices. Here, we analysed these two features on 32 matrices of quantitative pathogenicity trait data gathered from 15 plant-parasite pathosystems consisting of either annual or perennial plants along with fungi or oomycetes, bacteria, nematodes, insects and viruses. The performance of several nestedness and modularity algorithms was evaluated through a simulation approach, which helped interpretation of the results. We observed significant modularity in only six of the 32 matrices, with two or three modules detected. For three of these matrices, modules could be related to resistance quantitative trait loci present in the host. In contrast, we found high and significant nestedness in 30 of the 32 matrices. Nestedness was linked to other properties of plant-parasite interactions. First, pathogenicity trait values were explained in majority by a parasite strain effect and a plant accession effect, with no or minor parasite-plant interaction term. Second, correlations between the efficiency and scope of the resistance of plant genotypes, and between the host range breadth and pathogenicity level of parasite strains were overall positive. This latter result questions the efficiency of strategies based on the deployment of several genetically-differentiated cultivars of a given crop species in the case of quantitative plant immunity.

## Introduction

The effectiveness of strategies of disease control based on host immunity depends on the underlying capabilities of hosts to resist infection, of parasites to overcome this resistance and on the potential of these traits to evolve. Parasites and hosts can be specialists or generalists in, respectively, their capacity to infect and their immunity. Confronting multiple genotypes of a parasite with multiple genotypes of a host reveals their interaction patterns, *i*.*e*. the magnitude and arrangement of their mutual specialization or generalism, which gives insights into the underlying genetic bases of these characters and allows implementing strategies of disease management based on host diversification.

Importantly, the word “interaction” has different meanings in this context. In ecology, interactions between hosts and parasites are the effects that each of these two categories of living organisms have on each other. These host-parasite interactions can involve molecular interactions, which are attractive or repulsive forces between molecules, for example between parasite elicitors or effectors and host receptors. Finally, quantitative pathogenicity traits can be analysed thanks to statistical models that include, or not, a significant interaction between variables representing hosts and parasites. In the latter acception, “interaction” means that the model departs significantly from a purely additive model, including only a parasite effect and a host effect. Statistical interactions are used in the context of quantitative data and linear regression models, but not for qualitative binary data.

The structure of any host-parasite interaction can be represented as a matrix where columns correspond to host genotypes (either inbred lines, clones or F_1_ hybrids) and rows to parasite strains (either isolates, clones or populations depending on the considered parasite). Each cell in the matrix indicates the result of the pairwise confrontation between the corresponding host genotype and parasite strain. Qualitative interactions generate binary matrices with “1” and “0” grades, which correspond to successful and unsuccessful infections. These matrices are equivalent to networks, where links are represented between pairs of hosts and parasites that correspond to “1” grade in the matrix. Network theory has its origins in the study of social networks and in ecology of interacting organisms (Patterson and Atmar 1986). Ecological networks are typically identified by counting *in natura* the interactions between (or co-occurrence of) two sets of taxa. These analytical methods were recently used to analyse host-symbiont interactions resulting from cross-inoculation experiments, where every host taxon was inoculated with every symbiont taxon, and the compatibility of each host-symbiont pair was reported in the matrix (Flores et al. 2011; Flores et al. 2013; Weitz et al. 2013). The structural patterns of such matrices, where all host-symbiont pairs are evaluated under the same experimental and environmental conditions, are mainly the result of intrinsic, mostly genetic, differences between host or symbiont taxa.

Nestedness and modularity are two quantitative properties that reveal non-random distributions of “1” and “0” grades in such matrices or networks (Weitz et al. 2013). Nestedness measures the tendency of the hosts of a parasite to have a hierarchical organization, where the set of hosts of a given parasite (a species or a genotype) is a subset (respectively superset) of that of the parasites of broader (respectively narrower) host ranges. Here, the breadth of the host range of a given parasite is defined as the percentage of host species (or genotypes) that are infected by this parasite. The same tendency is observed for host immunity (Fig. 1A): the set of parasites that are controlled by the immunity of a given host is a subset (respectively superset) of that of hosts with broader (respectively narrower) scopes of resistance. Here, the scope of the resistance of a given host is defined as the percentage of parasite species (or strains) that are targeted by this resistance.

**Figure 1.**
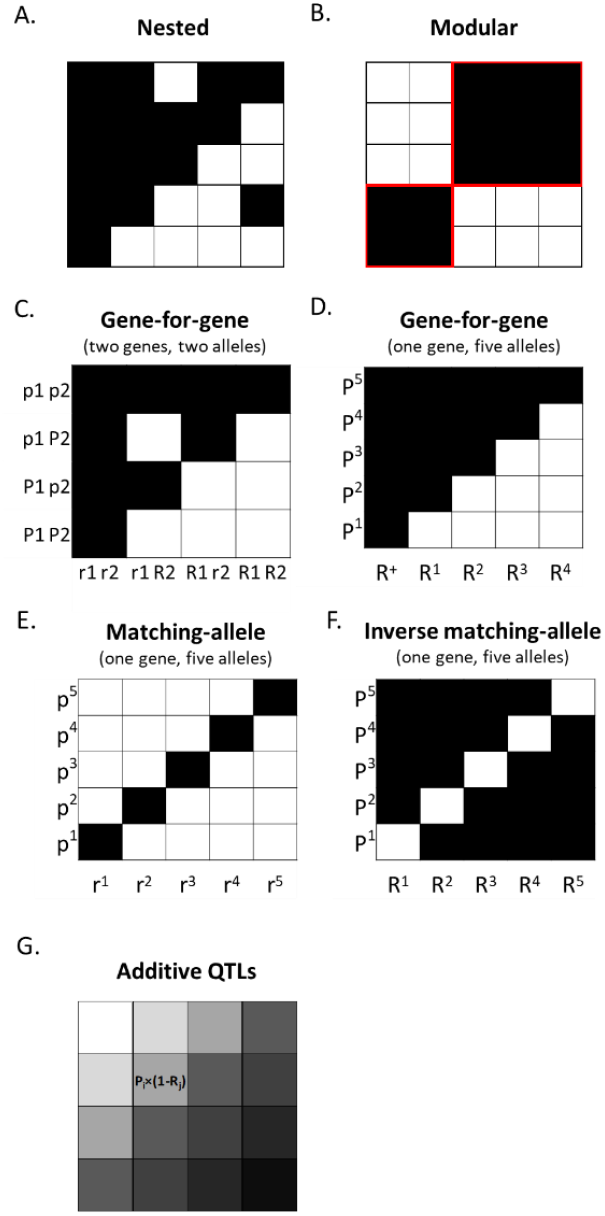
Matrices corresponding to different mechanistic, genetic and evolutionary models of qualitative or quantitative host-parasite interactions. In each case, host genotypes correspond to different columns and parasite genotypes to different rows) and black and white cells (or “1” and “0” grades) correspond to infection or lack of infection, respectively. A: Illustration of an imperfectly nested pattern. B: Illustration of a perfectly modular pattern (modules are delimitated with red lines). C and D: Gene-for-gene (GFG) models with partial or perfectly nested patterns. C: Case of two genes with two alleles in both hosts and parasites. Infection occurs only when no elicitor in the parasite is recognized by a product of the resistance alleles in the host. In the other situations, resistance is induced and there is no infection. D: Case of a single gene with five alleles in both hosts and parasites. Resistance alleles have various levels of specificity: in some plant accessions resistance can be induced by several parasite strains. E: Matching-allele model. Infection occurs only if the product of the pathogenicity allele is recognized by the product of the susceptibility allele in the host. F: Variation of D with higher specificity: resistance is induced by a specific product present in a single parasite genotype. This model was named “inverse matching-allele” model (Thrall et al. 2016) and has an anti-modular structural pattern. G: Additive QTL model with no plant-parasite QTL × QTL interaction. For each parasite strain i with pathogenicity level P_i_ and each plant accession j with resistance level R_j_, infection score corresponds to P_i_ x (1-R_j_).

Modularity measures the strength by which a matrix can be subdivided into a number of groups (*i*.*e*. modules) of hosts and parasites characterized by successful infections, while infections are rare between hosts and parasites that belong to different modules (Fig. 1B). Depending on the genetic, evolutionary and mechanistic patterns of host-parasite interactions, contrasted scores for nestedness and modularity are expected.

Evidence of nestedness is frequent for all kinds of matrices, including interactions between hosts and symbionts, either mutualistic or parasitic (Bascompte et al. 2003; Joppa et al. 2010; Dormann et al. 2017). A number of factors that are external to the interacting organisms can affect properties of such ecological networks. For example, nestedness increases with the abundance of taxa (Joppa et al. 2010; Staniczenko et al. 2013; Suweis et al. 2013; Valverde et al. 2018), with heterogeneous distribution of connections (*i*.*e*. numbers of links between interacting taxa; Jonhson et al. 2013), with the occurrence of broad connectivities (Feng and Takemoto 2014) and with spatially-limited interactions between taxa (Valverde et al. 2017).Three main models of host-parasite interactions have been proposed for qualitative plant-parasite interactions (Fig. 1C to F; see also Dybdahl et al. 2014 and Thrall et al. 2016). These models represent the mutual specialization of hosts and parasites in terms of underlying molecular mechanisms and genetic determinism and have consequences on the host-parasite coevolution pattern. Each model generates a specific structural pattern in the corresponding interaction matrix.

Historically, the first model was the gene-for-gene (GFG) model proposed to describe interactions between crop plants and their parasites (Flor 1956). In this model, immunity is induced upon recognition of a parasite elicitor by a host receptor, each encoded by a single gene. The loss or alteration of the elicitor in the parasite or the absence of a cognate resistance allele in its host results in infection. This model is coherent with dominant resistance that involves plant proteins containing nucleotide-binding and leucine-rich-repeat domains as receptors, and that mounts hypersensitive reactions (programmed cell death) upon recognition of parasite elicitors. The corresponding host-parasite matrix has a global nested pattern, with partial or complete overlap of the host ranges of the parasite strains and of the resistance spectra of the host genotypes (Fig. 1C,D; Gallet *et al*. 2016). Secondly, the matching-allele (MA) model was proposed to describe the self/non-self recognition system of invertebrate immunity (Grosberg and Hart 2000). In that case, infectivity requires a specific match between the host genotype and the parasite strain and, accordingly, universal infectivity is impossible. The corresponding host-parasite matrix has a modular structure. Cross-infections are frequent between hosts and parasites belonging to the same module but rare between hosts and parasites belonging to distinct modules. In extreme cases of specialization, modules can be as small as a single host-parasite pair (Fig. 1E). Mechanistically, this model is coherent with recessive plant resistance to viruses mediated by eukaryotic translation initiation factors (e.g. Sacristán and García-Arenal 2008) and with necrotrophic fungi which secrete elicitors of programmed cell death that increase plant susceptibility by allowing the fungus to feed on dying cells. In the context of plant necrotrophic parasites, this model is also confusingly named ‘inverse gene-for-gene’ (Peters et al. 2019). Thirdly, the inverse-matching-allele (IMA) model was proposed to reflect the adaptive immune system of vertebrates, where the host resists through recognition of the parasite and infections occur when the parasite mismatches the host (Kidner and Moritz 2013; Thrall et al. 2016). The IMA model was defined in the context of multi-allelic series of resistance and pathogenicity genes. Mechanistically very similar to the GFG model, it assumes that recognition between host and parasite genotypes is highly specific. The corresponding host-parasite matrix is therefore similar to the matching-allele model but with “0” and “1” grades replaced by “1” and “0” grades, respectively (Fig. 1F). Hence, a modular pattern is the expected result when immunity levels (instead of the degree of pathogenicity) are indicated in the matrix.

The distinguishing feature of the genetic models described above is that they describe qualitative binary interactions, where each host-parasite pair is characterized by its compatibility or non-compatibility. Models that describe quantitative host-parasite interactions are rare and their adequacy to represent empirical data have not been extensively tested (Lambrechts 2010; Boots et al. 2014; Wang et al. 2018). Analysis of quantitative plant immunity has mostly been confined to the framework of quantitative genetics and QTL (quantitative trait loci) mapping. These methods usually assume that resistance is determined by the additive effect of QTLs. More complex effects (dominance, epistasis) are rarely considered (Gallois et al. 2018). Furthermore, there are few studies of quantitative genetics and QTL mapping of parasite pathogenicity traits, especially in the case of plant parasites (Wang et al. 2018). Most importantly, these few analyses were conducted either with a set of hosts confronted to a single parasite or with a set of parasites confronted to a single host. In any case, there is a clear need for new models describing quantitative host-parasite interactions while properly accounting for the variability of both partners (Lambrechts 2010; Bartoli and Roux 2017). Moreover, previous work has shown that the outcome of analysis of matrix structure is markedly impacted when quantitative interactions are considered. Quantitative data are especially influencing the significance of nestedness (Staniczenko et al. 2013).

These considerations motivated us to conduct a comprehensive analysis of the nestedness and modularity of interaction matrices to deepen our knowledge in the specialization between plants and diverse parasites using quantitative data. The objectives of this work are (i) to assess the performance of available algorithms to identify nested and modular patterns in matrices of quantitative data and (ii) to determine if these patterns are specific to each pathosystem or show a general trend. In addition, our work provides a new perspective and insight into appropriate genetic and evolutionary models for representing quantitative plant-parasite interactions and for outcomes for plant resistance management.

## Results

We gathered 32 matrices corresponding to 15 plant-parasite pathosystems and containing quantitative pathogenicity trait values (Table 1; Fig. 2; Supplementary Methods 1). Among the 13 parasite species included, most were fungi or oomycetes (five and four, respectively), while bacteria, nematodes, insects and viruses were represented only once. Only three pathosystems included perennial (tree) plants and all plant species were temperate-climate crops (or crops adapted to both temperate and tropical climates). Each pathosystem included a set of strains belonging to the same parasite species and a set of accessions belonging to the same plant species with four exceptions, matrices 9, 18, 19 and 26, where accessions belonged to several closely-related plant species. Among the matrices, the number of plant accessions varied from seven to 53 (median 12) and the number of parasite strains varied from six to 98 (median 11.5). The number of matrix cells varied from 49 to 1470 (median 180). For most pathosystems, we analyzed several matrices corresponding to either different pathogenicity traits, different plant-parasite sets or different experiments. In order to meet the requirements of methods that allow the estimation of nestedness and modularity of matrices, the pathogenicity traits in each matrix were standardized into integer values ranging from 0 (minimal plant resistance and/or maximal parasite pathogenicity) to 9 (maximal plant infection and/or minimal parasite pathogenicity). We then tested for the occurrence of nestedness and modularity. For significance assessment, the nestedness/modularity scores of the matrices derived from experimental data were compared to those of simulated null-model matrices that are not expected to possess any nested or modular pattern (Supplementary Tables S1 to S18 in Supplementary Methods 2). Nestedness (or modularity) is significant if the actual matrix is more nested (or modular) than at least 95% of the matrices simulated under a given null model (black numbers on grey background in Tables 2 and 3 and in Supplementary Tables S19 and S20). As there are many possible null models and because their choice is crucial to conclude about the significance of nestedness or modularity, we analyzed the performance of the two available nestedness algorithms and of seven modularity algorithms in combination with different null models by estimating their type I and type II error rates through a simulation approach. A brief description of null models is provided in the Materials and Methods section. Details of null models and method performance analysis are provided in Supplementary Methods 2 and Supplementary Tables S1 to S18).

**Table 1:** Datasets used to analyze the structure of quantitative plant-parasite interaction matrices. AUDPC : Area under the disease progress curve.

| Matrix number | Parasite | Host plant | Matrix size (host × parasite) | Phenotype | Part of variance explained<sup>a</sup> | Reference or source |
| --- | --- | --- | --- | --- | --- | --- |
| **Bacterium** |  |  |  |  |  |  |
| 1 | *Pseudomonas syringae* | *Prunus armeniaca* (apricot) | 20 × 9 | Length of flat zone around the inoculation point at the surface of the shoot | 0.63 | Omrani *et al.*, unpublished |
| 2 | *P. syringae* | *P. armeniaca* | 20 × 9 | Length of flat zone around the inoculation point at the surface of the shoot | 0.81 | Omrani *et al.*, unpublished |
| 3 | *P. syringae* | *P. armeniaca* | 20 × 9 | Length of browning zone around the inoculation point below the bark of the shoot | 0.62 | Omrani *et al.*, unpublished |
| 4 | *P. syringae* | *P. armeniaca* | 20 × 9 | Length of browning zone around the inoculation point below the bark of the shoot | 0.70 | Omrani *et al.*, unpublished |
| **Fungi** |  |  |  |  |  |  |
| 5 | *Puccinia hordei* | *Hordeum vulgare* (barley) | 12 × 14 | Relative latent period | 0.81 | González *et al.*, 2012 |
| 6 | *Venturia inaequalis* | *Malus domestica* (apple) | 14 × 10 | % sporulating leaf area (AUDPC) | 0.57 | Laloi *et al.*, 2017 |
| 7 | *V. inaequalis* | *M. domestica* | 12 × 14 | % sporulating leaf area (AUDPC) | 0.40 | Caffier *et al.*, 2016 |
| 8 | *V. inaequalis* | *M. domestica* | 8 × 24 | % sporulating leaf area | 0.65 | Caffier *et al.*, 2014 |
| 9 | *Botrytis cinerea* | Tomato<sup>b</sup> | 12 × 94 | Lesion size on leaves | 0.55 | Soltis *et al.*, 2019 |
| 10 | *Podosphaera xanthii* | *Cucumis melo* (melon) | 19 × 26 | Sporulation surface on leaf disks | 0.92 | Dogimont *et al.*, unpublished |
| 11 | *P. xanthii* | *C. melo* | 19 × 31 | Sporulation surface on plants | 0.93 | Dogimont *et al.*, unpublished |
| 12 | *Zymoseptoria tritici* | *Triticum turgidum* subsp. *durum* (durum wheat) | 12 × 15 | % necrotic leaf area (AUDPC) | 0.73 | Marcel *et al.*, unpublished |
| 13 | *Z. tritici* | *T. turgidum* subsp. *durum* | 12 × 15 | % sporulating leaf area (AUDPC) | 0.72 | Marcel *et al.*, unpublished |
| 14 | *Z. tritici* | *Triticum aestivum* (bread wheat) | 15 × 98 | % necrotic leaf area | 0.59 | Marcel *et al.*, unpublished |
| 15 | *Z. tritici* | *T. aestivum* | 15 × 98 | % sporulating leaf area | 0.59 | Marcel *et al.*, unpublished |
| **Oomycetes** |  |  |  |  |  |  |
| 16 | *Phytophthora capsici* | *Capsicum annuum*<br>(pepper) | 10 × 6 | Necrosis length on stem (15 days post inoculation) | 0.98 | Messaouda *et al.*, 2015 |
| 17 | *P. capsici* | *C. annuum* | 53 × 6 | Necrosis length on stem (AUDPC) | 0.73 | Cantet *et al.*, unpublished |
| 17b<sup>c</sup> | *P. capsici* | *C. annuum* | 42 × 6 | Necrosis length on stem (AUDPC) | - | Cantet *et al.*, unpublished |
| 18 | *Phytophthora infestans* | *Solanum lycopersicum*<br>(tomato), *S. pimpinellifolium*, *S. habrochaites* and *S. pennellii* | 8 × 7 | Necrosis length on stem (AUDPC) | 0.86 | Ruellan *et al.*, unpublished |
| 19 | *Aphanomyces euteiches* | Fabaceae<sup>d</sup> | 8 × 35 | Root disease severity | 0.82 | Quillévéré-Hamard *et al.*, 2018 |
| 20 | *A. euteiches* | *Pisum sativum* | 10 × 43 | Root disease severity | 0.84 | Quillévéré-Hamard *et al.*, 2021 |
| 21 | *Plasmopara viticola* | *Vitis vinifera* (grapevine) | 8 × 33 | Necrosis on leaves | 0.72 | Paineau and Delmotte, unpublished |
| 22 | *P. viticola* | *V. vinifera* | 8 × 33 | Sporulation on leaves | 0.76 | Paineau and Delmotte, unpublished |
| **Insect** |  |  |  |  |  |  |
| 23 | *Aphis gossypii* | *C. melo* | 13 × 9 | Acceptance of plants | 0.52 | Boissot *et al.*, 2016 |
| 24 | *A. gossypii* | *C. melo* | 13 × 9 | Ability to colonize plants | 0.50 | Boissot *et al.*, 2016 |
| **Nematode** |  |  |  |  |  |  |
| 25 | *Globodera pallida* | *Solanum tuberosum*<br>(potato) | 10 × 20 | Cyst number (relative values) | 0.51 | Fournet *et al.*, unpublished |
| 26 | *G. pallida* | Wild potato species | 12 × 13 | Cyst eclosion rate | 0.78 | Gautier *et al.*, 2020 |
| **Virus** |  |  |  |  |  |  |
| 27 | Potato virus Y (PVY) | *C. annuum* | 7 × 8 | Virus load | 0.72 | Doumayrou *et al.*, unpublished |
| 28 | PVY | *C. annuum* | 7 × 9 | Virus load | 0.55 | Doumayrou *et al.*, unpublished |
| 29 | PVY | *C. annuum* | 7 × 8 | Symptom intensity (AUDPC) | 0.71 | Doumayrou *et al.*, unpublished |
| 30 | PVY | *C. annuum* | 7 × 7 | Symptom intensity (AUDPC) | 0.75 | Doumayrou *et al.*, unpublished |
| 31 | PVY | *C. annuum* | 7 × 8 | Relative dry matter weight | 0.55 | Doumayrou *et al.*, unpublished |
| 32 | PVY | *C. annuum* | 7 × 7 | Relative dry matter weight | 0.26 | Doumayrou *et al.*, unpublished |

<sup>a</sup> Unbiased estimator $\omega^2 = \omega^2_{\text{host}} + \omega^2_{\text{parasite}}$ , with $\omega^2_{\text{effect}} = (SS_{\text{effect}} - df_{\text{effect}} \times MS_{\text{error}}) / (SS_{\text{total}} + MS_{\text{error}})$ where SS is the sum of squares, df the number of degrees of freedom and MS the mean squares of the linear model: pathogenicity ~ 'parasite strain' + 'plant accession'.
<sup>b</sup> Two species: cultivated tomato (*Solanum lycopersicum*) and wild tomato (*S. pimpinellifolium*).
<sup>c</sup> Matrix 17b is identical to matrix 17 except that columns entirely made of zero-valued cells and redundant columns were removed.
<sup>d</sup> Four species: three pea (*Pisum sativum*) accessions, two vetch (*Vicia sativa*) accessions, two faba bean (*Vicia faba*) accessions and one alfalfa (*Medicago sativa*) accession.

**Figure 2.**
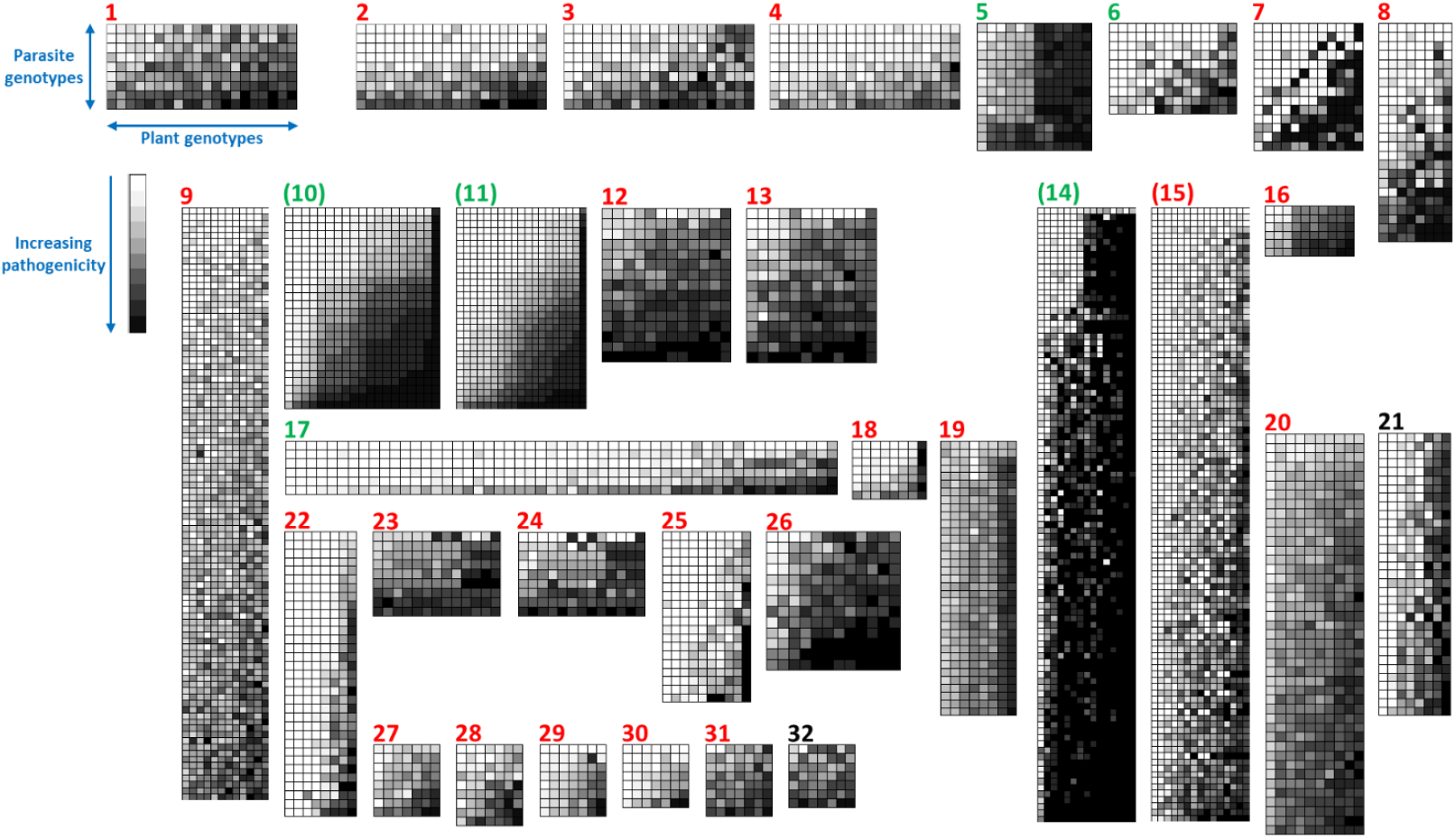
Overview of the 32 analyzed plant-parasite matrices (Table 1). Different plant accessions and parasite strains correspond to different columns and rows, respectively. White to black shades in each cell correspond to an increasing gradient of pathogenicity or infectivity (corresponding to 0 to 9 values in the analysed matrices) for a given plant and parasite pair. Rows and columns were ordered by increasing marginal totals, revealing the nested patterns. Colored numbers (red or green) correspond to significant nestedness (*WINE* algorithm) (Table 2). Green numbers correspond to significant modularity (*spinglass* algorithm), while numbers between parentheses correspond to significant modularity (*spinglass* algorithm) detected in reverse matrices.

**Table 2:** Analysis of nestnedness of plant-parasite interaction matrices with the *WINE* method.

| Matrix number | Nestedness score<sup>a</sup> | Null model<sup>b</sup> |  |  |  |  |  |  |
| --- | --- | --- | --- | --- | --- | --- | --- | --- |
|  |  | B | N | C1 | R1 | S | C2 | R2 |
| 1 | 0.78 | **0<sup>c</sup>** | 0 | 0 | 0 | 0 | 0 | 0 |
| 2 | 0.81 | **0.01** | 0 | 0 | 0 | 0 | 0 | 0 |
| 3 | 0.82 | 0 | 0 | 0 | 0 | 0 | 0 | 0 |
| 4 | 0.82 | 0 | 0 | 0 | 0 | 0 | 0 | 0 |
| 5 | 0.83 | 0 | 0 | 0 | 0 | 0 | 0 | 0 |
| 6 | 0.70 | 0.12 | 0 | 0 | 0 | 0 | 0 | 0 |
| 7 | 0.60 | 0.49 | 0 | 0 | 0 | 0 | 0 | 0 |
| 8 | 0.58 | **0.01** | 0 | 0 | 0 | 0 | 0 | 0 |
| 9 | 0.46 | 0 | 0 | 0 | 0 | 0 | 0 | 0 |
| 10 | 1.01 | 0 | 0 | 0 | 0 | 0 | 0 | 0 |
| 11 | 1.04 | 0 | 0 | 0 | 0 | 0 | 0 | 0 |
| 12 | 0.69 | 0 | 0 | 0 | 0 | 0 | 0 | 0 |
| 13 | 0.73 | 0 | 0 | 0 | 0 | 0 | 0 | 0 |
| 14 | 0.76 | 0 | 0 | 0 | 0 | 0 | 0 | 0 |
| 15 | 0.68 | 0.12 | 0 | 0 | 0 | 0 | 0 | 0 |
| 16 | 0.84 | 0 | 0 | 0 | 0 | 0 | 0 | 0 |
| 17 | 0.84 | 0 | 0 | 0 | 0 | 0 | 0 | 0 |
| 18 | 0.93 | 0 | 0 | 0 | 0 | 0 | 0 | 0 |
| 19 | 0.79 | 0 | 0 | 0 | 0 | 0 | 0 | 0 |
| 20 | 0.80 | 0 | 0 | 0 | 0 | 0 | 0 | 0 |
| 21 | 0.75 | 0.90 | 0 | 0.07 | 0 | 0 | 0.46 | 0 |
| 22 | 0.91 | 0 | 0 | 0 | 0 | 0 | 0 | 0 |
| 23 | 0.75 | 0 | 0 | 0 | 0 | 0 | 0 | **0.04** |
| 24 | 0.68 | 0.12 | 0 | 0 | **0.01** | 0 | 0 | **0.02** |
| 25 | 0.82 | **0.01** | 0 | 0 | 0 | 0 | 0 | 0 |
| 26 | 0.86 | 0 | 0 | 0 | 0 | 0 | 0 | 0 |
| 27 | 0.69 | 0.37 | 0 | 0 | 0 | 0 | 0 | 0 |
| 28 | 0.63 | 0.46 | 0 | **0.01** | 0 | 0 | 0 | 0 |
| 29 | 0.78 | **0.01** | 0 | 0 | 0 | 0 | 0 | 0 |
| 30 | 0.84 | **0.03** | 0 | 0 | 0 | 0 | 0 | 0 |
| 31 | 0.77 | 0 | 0 | 0 | 0 | 0 | 0 | **0.01** |
| 32 | 0.52 | 0.31 | 0.06 | 0 | 0.36 | 0.06 | **0.01** | 0.71 |

<sup>a</sup> Mean of 100 estimates.
<sup>b</sup> See Supplementary Methods 2 for details of the null models.
<sup>c</sup> Nestedness significance: the probability value (p-value) indicates the frequency of null-model matrices showing a strictly higher nestedness score than that of the actual matrix. P-values $\leq 0.05$ (significant nestedness) are in bold on grey cells and p-values $> 0.95$ (significant anti-nestedness) are in white on black cells.

**Table 3.**
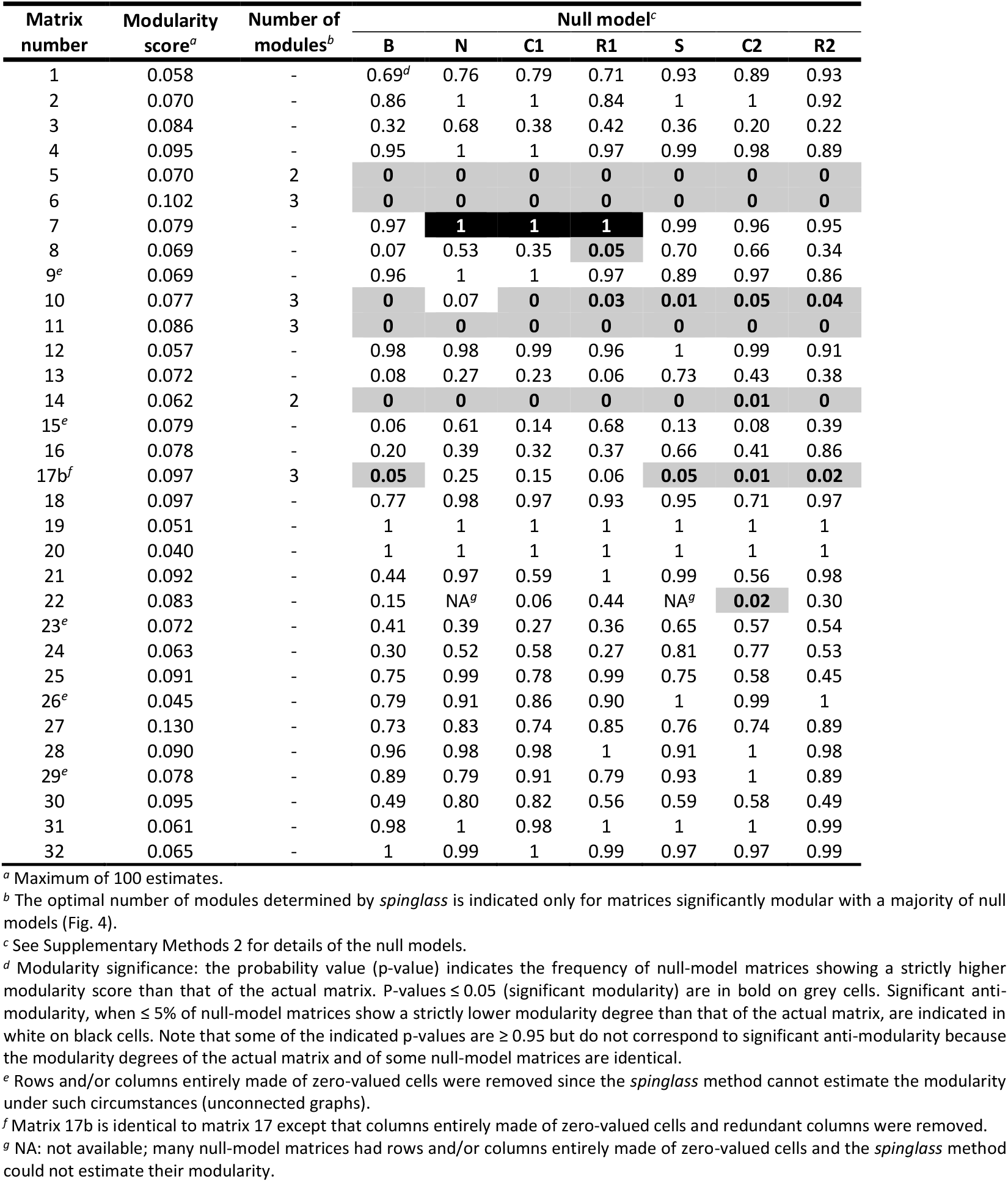
Analysis of modularity of plant-parasite interaction matrices with the *spinglass* method.

### Ubiquitous nestedness in quantitative plant-parasite interactions

First, we evaluated the performance of two algorithms, *WINE* and *wNODF* (Galeano et al. 2009; Almeida-Neto and Ulrich 2011), to estimate the nestedness of the 32 matrices. Simulations revealed that statistical significance with both null models C1 and R1 (or C2 and R2) provided the lowest false positive rates for nestedness (named null models CR1 or CR2; Supplementary Methods 2; Supplementary Tables S1 and S2). Moreover, the rates of false positive nestedness were similar for *WINE* and *wNODF* with null models CR1 or CR2. In contrast, a higher power was observed for *WINE* than *wNODF* for all null models (Supplementary Tables S5 and S6). Consequently, for the actual matrices, we focused mainly on the results of the *WINE* algorithm (Table 2).With the *WINE* algorithm, nestedness values were quite high in general (from 0.46 to 1.04; mean 0.77 on a scale varying from 0 to ≈1). The huge majority of the matrices (30/32; 94%) showed significant nestedness (p-values ≤ 0.05) with null models C1, R1, C2 and R2 (Table 2). Only matrices 21 and 32 were not significantly nested with either null model C1, R1, C2 or R2.

Fewer matrices showed significant nestedness with the *wNODF* algorithm (Supplementary Table S19), which is consistent with the lower statistical power of *wNODF* compared to *WINE*.

### Investigation of the biological significance of nestedness

#### Adequacy of an additive linear regression model for pathogenicity matrices

The high and significant nestedness observed among most of the analysed matrices suggests that an additive model combining pathogenicity QTLs in the parasites and resistance QTLs in the hosts, but omitting QTL x QTL interactions between hosts and parasites, would fit well with the data (Fig. 1G). We evaluated the performance of the linear regression model: ‘pathogenicity’ ∼ ‘parasite strain’ + ‘plant accession’, with no interaction term, on the datasets. For each plant accession-pathogen strain pair, the mean pathogenicity value was considered for the ‘pathogenicity’ variable. The ‘parasite strain’ and ‘plant accession’ effects were highly significant (p-value < 0.0012), except for matrices 21 and 32 which were the only ones not significantly nested according to the *WINE* method (Table 2). Omitting these two matrices, the unbiased estimate of the part of variance explained by the ‘parasite strain’ and ‘plant accession’ effects (ω^2^; Kirk 1982) varied from 0.40 to 0.98 (mean 0.69) (Table 1), which lends support to the suggested genetic model. Only one ω^2^ value was below 0.50 (matrix 7). Moreover, the ω^2^ values of the linear regression model were significantly correlated with the nestedness scores obtained with the *WINE* algorithm (Pearson’s r = 0.74; p-value = 1.5e-06) across the 32 matrices. They were only marginally correlated with the nestedness scores of the *wNODF* algorithm (r = 0.31; p-value = 0.087).

To confirm these results, we estimated the part of phenotypic variance explained by the ‘parasite × plant’ interaction effects for the different matrices using the linear regression model: ‘pathogenicity’ ∼ ‘parasite strain’ + ‘plant accession’ + ‘parasite strain × plant accession’. To disentangle the interaction effect from the uncontrolled environmental and experimental variance, several independent phenotypic values should be available for each plant-parasite genotype pair. Such values are not available for all published data and for some of our experimental data. Consequently, we obtained estimates of the parasite × plant interaction for 26 of the 32 matrices and Soltis et al. (2019) provided an estimate for a 27^th^ matrix (Supplementary Table S20). For five of these matrices (1, 2, 9, 12 and 13), the parasite × plant interaction was not significant (Supplementary Table S20 and Soltis et al. 2019). In the 24 significantly-nested matrices among the 26 analyzed, the plant-parasite interaction term explained on average ω^2^ = 17.5% (minimum 0% ; maximum 41%) of the phenotypic variance, relatively to the total phenotypic variance that could be collectively explained by the plant, the parasite and the plant-parasite effects. The average ω^2^ was 18.9% for the 27 analyzed matrices.

Finally, for matrices 5, 10, 11 and 16, for which no estimate of the plant-parasite interaction effect could be obtained, the plant and parasite effects alone explained 81%, 92%, 93% and 98% of the phenotypic variance, respectively (Table 1). Hence, a minor interaction effect, if any, is also expected for these latter four matrices. For a single significantly-nested matrix (number 7), the plant and parasite effects alone explained a minor part of the phenotypic variance (40%; Table 1) and we were unable to obtain an estimate of the plant-parasite interaction effect.

Collectively, these results support the fact that the parasite × plant interaction determines a minor part, if any, of the variation of quantitative pathogenicity traits.

#### Evaluating potential trade-offs: Host range breadth vs. pathogenicity in parasites and scope vs. efficiency of resistance in host plants

The ubiquitous nestedness detected suggests a positive correlation between the host range breadth, *i*.*e*. the percentage of host accessions that a parasite can efficiently infect, and the pathogenicity level of the parasite. Similarly, a positive correlation is expected between the scope of the resistance and the resistance efficiency of the plants. Given the continuous distribution of the quantitative pathogenicity traits, we defined arbitrary pathogenicity thresholds to distinguish host and non-host accessions for a given parasite strain, and to distinguish parasite strains included or not included in the scope of the resistance of a given plant accession. Nine thresholds were defined, varying from 10% to 90% of the maximal pathogenicity value in the whole matrix by increments of 10%, and allowed estimating the percentage of plant accessions included in the host range of each parasite strain (*i*.*e*. the host range breadth) and the percentage of parasite strains included in the scope of resistance of each plant accession. The mean Pearson’s coefficient of correlation (r) between host range breadth and pathogenicity varied from 0.20 to 0.38 across the different threshold values (mean 0.31). Depending on the threshold, from 23.1% (6/26 matrices) to 40.6% (13/32) (mean 31.9%) of the matrices showed significantly positive r values, whereas from 0 (0/11) to 9.7% (3/31) (mean 4.8%) of the matrices showed significantly negative r values (Fig. 3; Supplementary Table S21). Note that the coefficient of correlation could not be calculated for several matrices for some of the thresholds because of the lack of pathogenicity values above (for correlation between host range breadth and pathogenicity) or below (for correlation between resistance scope and efficiency) that threshold. The mean r between resistance scope and efficiency varied from 0.18 to 0.59 across the different threshold values (mean 0.39). Depending on the threshold, from 25.0% (6/24) to 48.4% (15/31) (mean 36.6%) of the matrices showed significantly positive r values, whereas from 0 (0/32) to 9.4% (3/32) (mean 3.0%) of the matrices showed significantly negative r values (Fig. 3).

**Figure 3.**
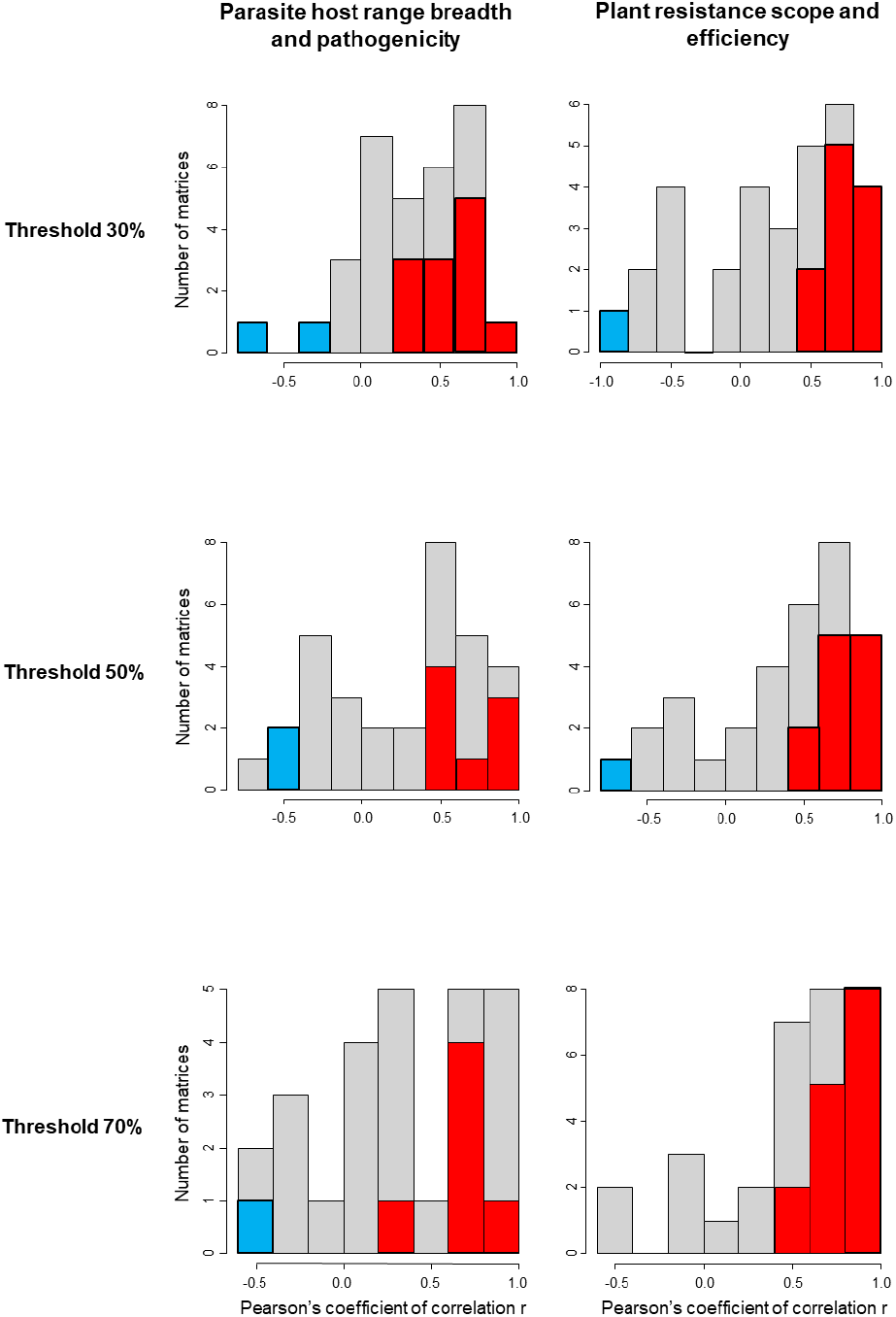
Distributions of Pearson’s coefficients of correlation (r) between parasites host range breadth and pathogenicity (left) or between plant resistance efficiency and scope (right) across the 32 analysed matrices for different thresholds separating hosts and non-hosts (or parasites included or not included in the resistance scope). Each threshold corresponds to a percentage of the maximal pathogenicity value in each matrix (only results obtained with thresholds corresponding to 30%, 50% and 70% of the maximal pathogenicity value are shown; results were similar for other thresholds). In blue and red: significantly negative or positive r values (p-value < 0.05). For some thresholds and some matrices, the coefficient of correlation could not be calculated because too few pathogenicity data remained.

### Rare cases of modularity in quantitative plant-parasite interactions

As for nestedness, thanks to a simulation approach we could evaluate the performance of four of the modularity algorithms: *edge betweenness* (Newman and Girvan 2004), *fast greedy* (Clauset et al. 2004), *louvain* (Blondel et al. 2008) and *spinglass* (Newman and Girvan 2004; Reichardt and Bornholdt 2006; Traag and Bruggeman 2009) (Supplementary Methods S2). By maximizing a modularity score, these algorithms estimate the optimal number of modules and the distribution of plant and parasite genotypes in the modules. In terms of type I error rate, the *spinglass* method was by far the most efficient for all null models (Supplementary Tables S7 to S10). For the other three methods, the type I error rate varied greatly depending on the null models, S, C2, R2 and CR2 being the most efficient. The *spinglass* method had also the lowest false positive rate to detect anti-modularity, i.e. a situation where a matrix is less modular than at least 95% of the matrices simulated under a given null model (Supplementary Tables S11 to S14). Consequently, we focused on the *spinglass* method to detect modularity in the 32 actual matrices (Table 3).

Modularity scores were low overall, with a maximum of 0.130 and a mean of 0.077, on a scale varying from 0 to 1 (Table 3). Six matrices (numbers 5, 6, 10, 11, 14 and 17b) were significantly modular with a majority of null models (Table 3), though their modularity scores were low (≤ 0.102). Depending on the matrix, *spinglass* defined an optimal number of two or three modules, which provided the maximal modularity score (Table 3; Fig. 4). In addition, matrices 8 and 22 were only significantly modular with one null model.

**Figure 4.**
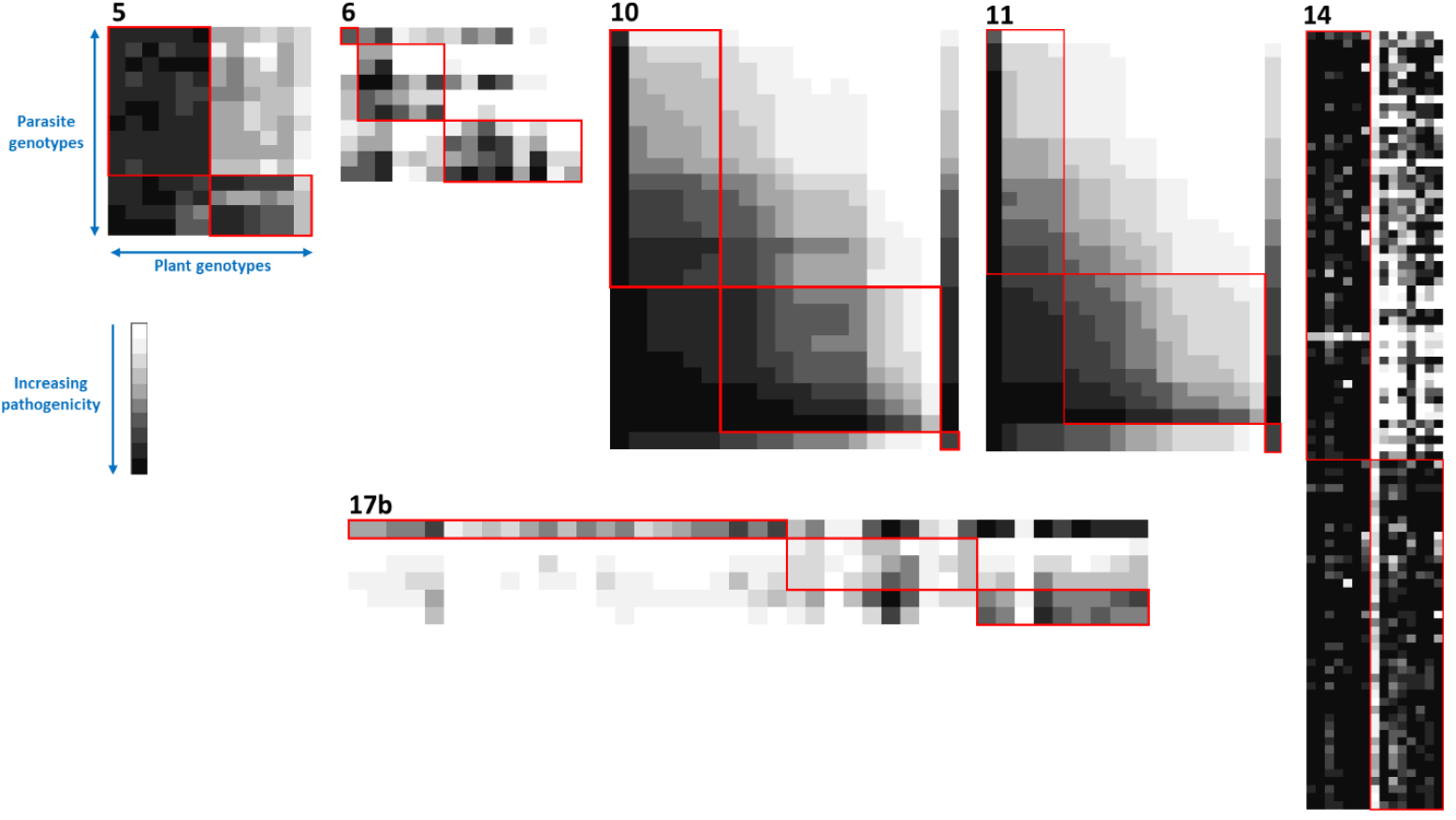
Overview of the six plant-parasite matrices showing significant modularity with the *spinglass* algorithm (Table 3). Rows and columns were ordered by modules, delimited by red lines. See legend of Fig. 2 for the representation of matrices.

The *edge betweenness, fast greedy* and *louvain* methods did not allow to detect consistently significant modularity in any matrix (Supplementary Table S22).

The *spinglass* algorithm showed also that matrix 7 was significantly anti-modular with null models N, C1 and R1 (Table 3). The *edge betweenness, fast greedy* and *louvain* methods detected significant anti-modularity in most matrices with most null models (Supplementary Table S22) but suffered high rates of false positive anti-modularity for many null models (Supplementary Methods 2; Supplementary Tables S11 to S13).

### Investigation of the biological significance of modularity

We examined the relevance of the detected modules for the six matrices showing significant modularity with most null models with *spinglass* (Table 2) by analysing whether the plant and parasite genotypes belonging to each module shared common properties (common resistance gene or QTL for plants; common pathogenicity factor for parasites; common origin for plants or parasites).

For matrix 5 (*Puccinia hordei*-barley), two modules were detected (Fig. 4). The first one grouped the five accessions with resistance QTLs *Rphq3* and *Rphq11*, showing delayed infection with most isolates of the second module, and one accession carrying QTLs *Rphq1, Rphq2* and *Rphq3*, showing delayed infection with almost all isolates (González et al. 2012). The second module contained four accessions with either no resistance QTL or QTL *Rphq18*, that were quickly infected by almost all isolates. The country of origin or date of collection of the isolates did not explain their distribution in the two modules (Marcel et al., 2008).

For matrix 6 (*Venturia inaequalis*-apple), three modules were detected. The first one grouped the eight accessions carrying QTL *T1* and the four *V. inaequalis* isolates collected on apple trees carrying *T1* (Laloi et al. 2017). The two other modules grouped (i) the remaining accessions that were either carrying no resistance QTL or QTLs F11 or F17 that have only a low effect on disease reduction and (ii) isolates collected on these accessions. One of these modules grouped a single isolate and a single accession. Infections were on average high within all modules and low between any pair of modules.

Two modules were also detected for matrix 14 (*Zymoseptoria tritici*-bread wheat). These modules could be partially explained by the interaction between the resistance gene *Stb6* (Saintenac et al. 2018), that confers a high level of resistance in the absence of a hypersensitive response, and the pathogen avirulence gene *AvrStb6* (Zhong et al. 2017). Six of the eight cultivars in the first module carry *Stb6*, while at least six of the seven cultivars in the second module do not carry *Stb6*. Moreover, the 44 fungal isolates structuring the first module are pathogenic on *Stb6* while the 54 isolates from the second module are either pathogenic or not pathogenic on *Stb6*.

Concerning matrices 10, 11 (*Podosphaera xanthii*-melon) and 17b (*Phytophthora capsici*-pepper), three modules were detected but there was no evidence of similarity in the genetic composition of accessions, the presence of particular resistance genes or QTLs or the origin of isolates belonging to a same module.

### Modularity of reverse matrices

To test the occurrence of IMA patterns (Fig. 1F), we also analyzed the modularity of the 32 matrices transformed such that “0” grade corresponds to the maximal plant susceptibility and “1” to “9” grades correspond to the range of increasing plant resistance (hereafter “reverse matrices”). Using the *spinglass* algorithm, four matrices (numbers 10, 11, 14 and 15) showed significant but low modularity (≤0.078) with either null models C1 and R1 or C2 and R2. Depending on the matrix, *spinglass* defined an optimal number of two to five modules (Fig. 5).

**Figure 5.**
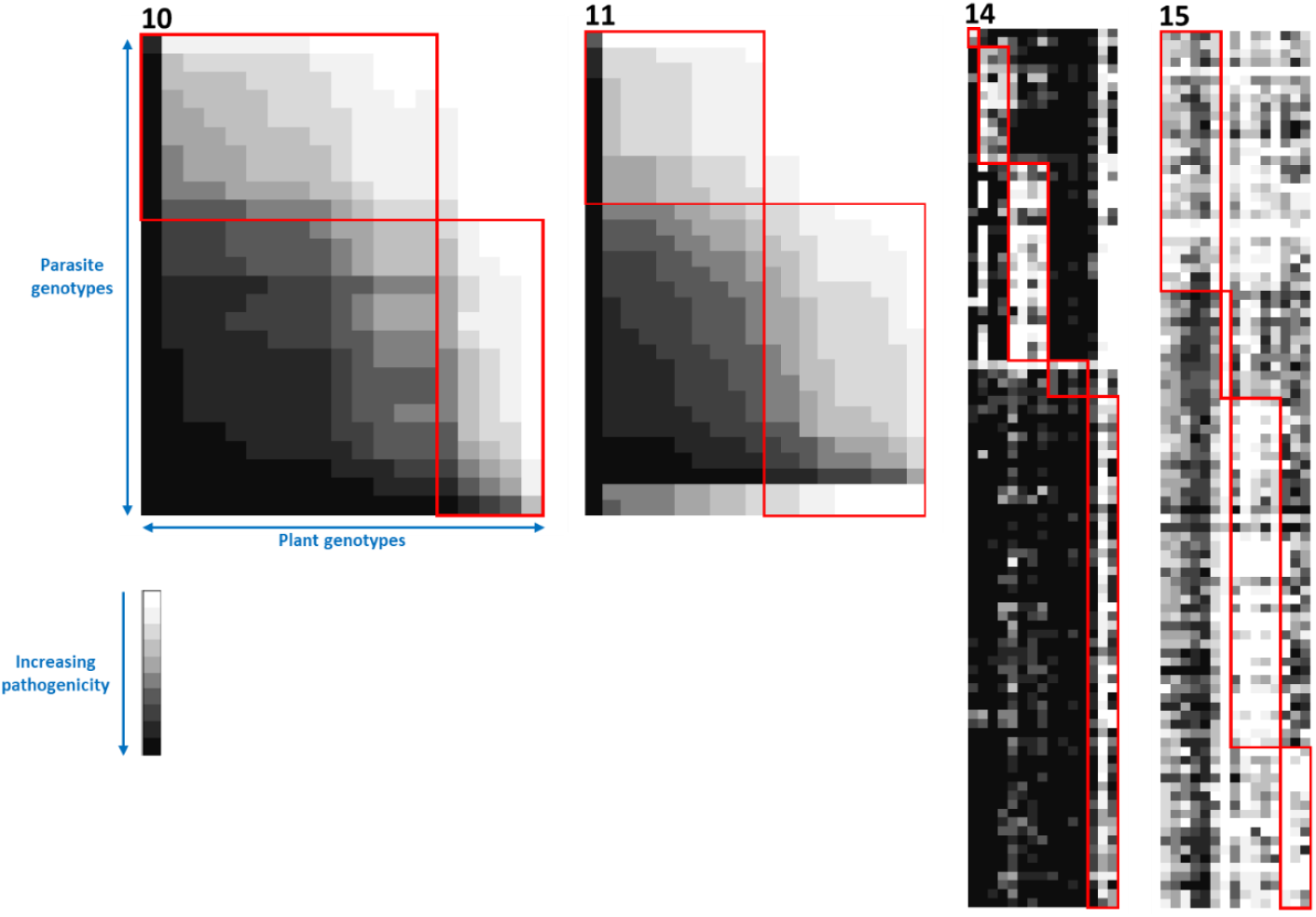
Overview of the four plant-parasite matrices showing significant modularity with the *spinglass* algorithm when matrices were transformed such that 0 values correspond to the maximal plant susceptibility and 9 values to the maximal plant resistance (but note that the matrices are represented such that 0 to 9 values correspond to a plant resistance to susceptibility gradient, as in the original matrices). Rows and columns were ordered by modules, delimited by red lines. See legend of Fig. 2 for the representation of matrices.

The modules identified in reverse matrices 14 and 15 using the *spinglass* algorithm were biologically more meaningfull than the two modules previously identified for matrix 14. Matrices 14 and 15 correspond to two different phenotypic traits measured in the same plant-parasite interactions (i.e. necrosis and sporulation, respectively). Interestingly, modules identified in the two matrices were similar but not identical since five modules were identified in matrix 14 and four modules were identified in matrix 15. This may reflect differences in the genetic determinism of the two phenotypic traits measured or differences in the mechanisms of various *Stb* resistance genes. For matrix 14, three modules correspond to the presence of resistance genes *Stb7* (one cultivar), *Stb9* (three cultivars) and *Stb6* (four cultivars), one module to cultivars carrying various *Stb* genes (three cultivars), and one module to susceptible (or partially resistant) cultivars (four cultivars). For matrix 15, the modules corresponding to the presence of *Stb6* and *Stb9* are also identified (with an additional cultivar in the *Stb6* module), the module corresponding to susceptible cultivars as well (with two additional cultivars), and the cultivar Salamouni carrying *Stb13* and *Stb14* forms the fourth module. As above, there was no evidence of similarity in the composition of accessions and isolates belonging to the same module for reverse matrices 10 and 11.

Overall, considering both the initial and reverse matrices, our analysis revealed that only a minority of the matrices (7/32; 22%) were significantly modular. When several matrices were analyzed for the same patho-system, some discrepancies could be observed between matrices. For nestedness, matrix 21 was not significantly nested, whereas matrix 22 was. Similarly, among matrices 27 to 32, only matrix 32 was not significantly nested. Similar observations could be made for modularity analyses. These differences could be due to the specific pathogenicity trait that vary between matrices and/or to the statistical power to detect the nested or modular structures in these matrices.

## Discussion

There is nothing more fundamental to the concepts in Plant Pathology as a science and to the practical strategies used for managing plant health than the host range of a parasite and the scope of resistance of a plant (Morris and Moury 2019). Based on the patterns in matrices of plant-parasite interactions, we can conceive and test hypotheses about the molecular and evolutionary processes that underlie plant-parasite interactions, develop robust diagnostic tools, design breeding programs and strategies for deploying resistant cultivars, and construct models to anticipate disease emergence. Given the complexity of the mechanisms involved in disease, it would be reasonable to assume that the particularities of each pathosystem would be an impediment to identifying universal principles that can guide these efforts. However, here we have used network-based analyses to reveal the quasi-universal principle that the structure of quantitative matrices of plant-parasite interactions is nested. Indeed, evidence of nestedness was found in 94% (30/32) of the matrices that we analyzed. Our results were based on statistically robust analyses of quantitative assessments of compatible interactions between hosts and parasites for large interaction matrices involving from 49 to 1470 (median 180) host-parasite combinations. Quantitative data are key to the accuracy and genericity of these analytical methods. Indeed, in a study of 52 published matrices containing data on plant-pollinator, plant-seed disperser and parasitoid-host interactions, Staniczenko et al. (2013) found evidence of nestedness in only 3% of matrices including quantitative data, whereas the same matrices considered in a binary manner showed evidence of nestedness in 98% of cases.

Network analyses can also be strongly affected by the choice of null models (Gotelli and Graves 1996). This is why we conducted a thorough evaluation of the performance of several null models with simulations (Supplementary Methods 2). The null models should keep, as much as possible, everything identical to the actual matrix apart from the pattern of interest, nestedness or modularity. Many null models have unacceptably loose constraints. For example, null models that do not force row or column marginal sums to be constant create distributions of taxa that do not match those usually observed, leading to falsely positive nestedness (Brualdi and Sanderson 1999; Joppa et al. 2010). Accordingly, high rates of false positives were observed with null models N and S in our simulations (Supplementary Tables S1 and S2). Since parasites typically differ greatly in the number of hosts they exploit and the efficiency with which they exploit them, we did not want null models to detect significant nestedness when the heterogeneity of infection was shuffled randomly among hosts, as was frequently observed for null models N and S with test matrices M1R to M5R (Supplementary Tables S1 and S2). Null models R1 and R2 that force row marginal sums to be constant avoided this problem (Supplementary Tables S1 and S2). The same was true for the scope and efficiency of resistance that differ greatly between plant accessions. In that case, the C1 and C2 null models efficiently avoided an excess of falsely positive nestedness due to the hererogeneity of resistance (because C1 and C2 are equivalent to R1 and R2 when the rows and columns of the matrix are exchanged, which leaves the nestedness scores unchanged; data not shown). Overall, to account for both plant resistance and parasite infection heterogeneities, we found that the CR1 (or CR2) null model, that combines null models C1 and R1 (or C2 and R2, respectively), is the most efficient as it showed acceptable type I error rates (Supplementary Methods 2). Null model B, based on Patefield’s (1981) algorithm, maintains both the row and column marginal sums of the actual matrix. However, it does not maintain the connectance (*i*.*e*. number of non-zero-valued cells of the matrix), which has a strong impact on the estimation of nestedness. Consequently, the type I error rates associated with null model B were frequently higher than those obtained with models CR1 or CR2. Moreover, using quantitative instead of binary data contributed to lowering the nestedness false positive rate (Staniczenko et al. 2013; Dormann et al. 2017).

Overall, we obtained strong and consistent evidence of nestedness for almost all matrices (except matrices 21 and 32), whatever the parasite type, the plant species or the pathogenicity trait measured. In this analysis, we analysed essentially crop plant systems. The same kind of nested structure could be expected in wild plant-parasite systems, since quantitative interactions are probably more frequent in natural pathosystems compared to crop plants, where major-effect resistance genes have been primarily bred (Boots et al. 2014). Nestedness was linked to two important features of quantitative plant-parasite matrices: (i) low level of statistical interactions between plant and parasite genotypes in terms of infection intensity and (ii) lack of trade-offs between host range and pathogenicity among parasite strains and between efficiency and scope of the resistance among plant accessions.

The low level of plant-parasite statistical interaction is supported by the fact that an additive linear model - containing only a plant accession effect and a parasite strain effect with no interaction term - explained a high proportion of the phenotypic variance, ω^2^ varying from 0.40 to 0.98 across matrices (mean = 0.69) (Table 1). In addition, we could estimate the plant-parasite interaction effect for 27 of the 32 matrices (Supplementary Table S20). The interaction was not significant for five matrices and it explained, on average, only 11.8% of the total trait variance or only 18.9% of the variance that could be explained collectively by the plant, the parasite and their interaction effects.

This result is compatible with a genetic model where pathogenicity in the parasite and resistance in the host plant are determined by a varying number of QTLs, but the statistical interaction between effects of QTLs from the parasite and QTLs from the host is rare and/or of small magnitude (Table 1; Supplementary Table S20; Fig. 1G). In other words, plants and parasites differ by their QTL assemblage (i.e. QTL numbers and/or effects) but plant resistance QTLs have similar effects towards all parasite strains and, reciprocally, parasite pathogenicity QTLs have similar effects towards all plant genotypes. This model is similar to the one proposed in Fig. 1A by Boots *et al*. (2014), who explored its consequences in terms of host-parasite coevolution. Quantitative models usually used to analyse empirical data on plant-parasite interactions are quite simplistic,

*e*.*g*. assuming or not a statistical interaction between plant and parasite genotypes (Parlevliet 1977). Models that are more complex have been proposed in the frame of theoretical modelling (*e*.*g*. Sasaki 2000; Fenton *et al*. 2009; Fenton *et al*. 2012; Boots *et al*. 2014) but their relevance to represent biological data was not evaluated. Importantly, we do not argue that evidence of nestedness supports a single genetic model of plant-parasite interaction. Instead, we suggest that an additive linear model with a plant accession and a parasite strain effects is the simplest model that accounts for the empirical data but other models could be suitable, like the modified GFG models of Sasaki (2000), Fenton *et al*. (2009) or Boots *et al*. (2014). A future challenge, requiring more in-depth genetic studies, would be to evaluate the adequacy of these different models to represent empirical plant-parasite interactions. New analytical methods can provide a better understanding and quantification of host-parasite genetic interactions, such as the host-parasite joint genome-wide association analysis recently developed by Wang et al. (2018). Applied to the *Arabidopsis thaliana*– *Xanthomonas arboricola* pathosystem, this model showed that 44%, 2% and 5% of the phenotypic variance could be explained respectively by the parasite strain, the host accession and the parasite-host interaction. As in our results, only a small parasite-host interaction effect was detected.

Models of host-parasite interaction, including a few quantitative ones, were extensively used in theoretical modelling to analyse their consequences in terms of host-parasite coevolution, a situation that is more appropriate to wild pathosystems than to crops, where the host plants are not allowed to evolve freely but are chosen by growers. One widely explored question is how the host and parasite diversities are maintained along time. In the case of GFG models or quantitative models with no host-parasite interactions, costs to resistance in the hosts and infectivity in the parasite are critical to the static polymorphism and to the set up of coevolutionary dynamics in host and parasite populations (Sasaki 2000; Brown and Tellier 2011; Fenton *et al*. 2012; Boots et al. 2014). However, we did not observe costs to the scope of resistance in the plants or to the host range breadth in the parasites in the present study, but instead globally positive relationships with resistance strength and parasite pathogenicity, respectively (see below).

The lack of trade-offs in plants or parasites is supported by the fact that we observed a majority of positive, rather than negative correlations (i.e. trade-offs), between the infectivity and the breadth of host range of parasites on the one hand and, especially, between the efficiency and scope of the resistance of plants on the other hand (Fig. 3). Few studies have examined the relationships between the scope and efficiency of plant resistance. In contrast with our results, Barrett et al. (2015) hypothesized evolutionary trade-offs between resistance efficiency and scope because quantitative resistance had a broader scope compared to qualitative resistance in the *Linum marginale* – *Melampsora lini* interactions. The difference between our studies could be that we focussed on quantitative resistance and included few qualitative resistance genes in our dataset (or these were overcome by most parasite strains). The positive correlation between parasite infectivity and host range breadth contrasts with qualitative host-parasite interactions and especially the GFG model, where the expansion of the host range of parasites is associated with a cost in fitness during infection of the previous hosts. Such so-called “virulence costs” have been experimentally measured in many plant-parasite systems, including viruses (Jenner et al. 2002; Desbiez et al. 2003; Janzac et al. 2010; Poulicard et al. 2010; Fraile et al. 2011; Ishibashi et al. 2012; Khatabi et al. 2013), fungi (Bahri et al. 2009; Huang et al. 2010; Caffier et al. 2010; Bruns et al. 2014), oomycetes (Montarry et al. 2010), bacteria (Vera Cruz et al. 2000; Leach et al. 2001; Wichmann and Bergelson 2004) or nematodes (Castagnone-Sereno et al. 2007), and could explain why universal pathogenicity is not fixed in pathogen populations (Tellier and Brown 2011). For quantitative plant resistance, few studies have estimated the occurrence of pathogenicity costs. Montarry et al. (2012) showed a cost for PVY to adapt to a quantitative pepper resistance when inoculated to a susceptible pepper genotype, whereas Delmas et al. (2016) showed, on the opposite, that there was no fitness cost associated with the adaptation of *Plasmopara viticola* to partially resistant grapevine varieties. Fournet et al. (2016) even highlighted that nematode populations that had adapted to potato quantitative resistance were more pathogenic on a susceptible potato genotype than were naïve nematode populations. The present study focused mostly on interactions between plants and parasites at the intraspecific level, but other studies have revealed a similar trend when strains of a given parasite species are confronted with numerous plant species. For example, a positive correlation was observed between species host range and pathogenicity for *Pseudomonas syringae* (Morris et al. 2000, 2019). For this bacterium, the most pathogenic strains were also the most ubiquitous in the environment, suggesting also an absence of trade-off between host range and dispersal capability or survival in the environment (Morris et al. 2010).

In contrast to nestedness, we obtained little evidence of modularity among the matrices that we analysed. Modularity scores were low for all matrices. In only seven matrices, representing either infection or resistance scores (i.e. reverse matrices), did we detect significant modularity with a majority of null models (Tables 3 and 4; Fig. 4 and 5). For four of these matrices (matrices 5, 6, 14 and 15), modularity was linked to the presence of particular resistance genes or QTLs in the plant accessions and, for the parasite strains, to the presence of particular avirulence genes or to a common origin in terms of host genotype. For the remaining matrices (10, 11 and 17b), no common property could be found for plant accessions and parasite strains belonging to the same module. The lack of modularity of infection matrices and of reverse matrices suggests that the MA and IMA genetic models are either inadequate to represent the structure of quantitative plant-parasite interactions or explain only marginally their structure (Fig. 1E,F).

## Conclusion

The ubiquitous nested patterns observed in quantitative plant-parasite interaction matrices have important implications for our understanding and management of plant diseases. They can help infer the underlying genetic bases of quantitative aspects of disease manifestation and their evolution. Our results are compatible with an additive model comprising a plant resistance effect, a parasite pathogenicity effect and no (or little) plant-parasite interaction effect. Given the relatively small number of pathosystems (15) analysed here, it is not yet possible to assess if these patterns differ according to the parasite type (viruses, bacteria etc…; obligate or facultative), the plant type (perennial or annual; crops or wild). Obtaining experimental data from additional cross-inoculation experiments and analysing the structure of the resulting matrices could help answer these questions.

A major enigma that we highlight is the apparent lack of trade-off between pathogenicity and host range breadth among strains of a parasite, which has important implications on the efficiency of plant resistance management through cultivar rotation, mixtures or mosaics. Indeed, these strategies rely at least in part on a counter-selection of the most pathogenic parasite strains by a diversification of plant cultivars (Brown 2015). The efficiency of these strategies would certainly be reduced in absence of costs of adaptation to plant resistance. Therefore, in absence of such costs, the efficiency of the rotation, mixtures or mosaic strategies would rather depend on barrier effects, *i*.*e*. effects of plants that hamper the dispersal of parasites in agricultural landscapes through their architecture or through repellent volatile organic compounds.

## Materials and Methods

### Datasets

To be able to analyse plant-parasite interaction networks, we selected published or unpublished datasets containing at least 6 plant accessions and 6 parasite strains. A brief description of these datasets is provided in Table 1. A more exhaustive description of these datasets (characteristics of plant accessions, parasite strains and phenotyping procedures) is provided in Supplementary Methods 1.

### Network analyses

The nestedness and modularity of the different matrices were estimated, and their statistical significance tested respectively with the ‘bipartite’ and ‘igraph’ packages of the R software version 3.5.1 (http://cran.r-project.org/). These analyses were initially developed for the study of social, then of ecological, networks (or equivalently matrices) containing counts of links between individuals or between interacting species. Hence, to perform these analyses, the matrices should only contain integer values. Moreover, some nestedness or modularity algorithms cannot run in the absence of zero-valued matrix cells or in the presence of an excess of zero-valued cells leading to an unconnected network.

Consequently, the first step consisted in transforming the actual matrices accordingly. In all matrices, pathogenicity trait values were transformed into integers from 0 to 9. For this, ten intervals with equal sizes and spanning the range of the pathogenicity trait values of the actual matrix were defined. The bounds of these intervals are [P_min_ + (P_max_ – P_min_)\**i*/10, P_min_ + (P_max_ – P_min_)*(*i* + 1)/10], with *i* being an integer in the [0,9] interval and P_max_ and P_min_ being the maximal and minimal pathogenicity trait values in the whole matrix, respectively. Then, depending on its inclusion in a given pathogenicity trait value interval defined as above, each matrix value was transformed into the corresponding *i* integer value. When necessary, the matrix was modified in order that “0” and “9” grades correspond to the minimal and maximal pathogenicity classes, respectively, and not the opposite. A continuous distribution of the pathogenicity grades was observed in 30 of 32 matrices (Fig. 2). However, for matrices 17b and 22 that contained a large number of zero-values cells, phenotypic values were log-transformed to spread out the data more evenly among the ten phenotypic classes. As these log-transformed matrices produced similar results as the actual matrices in terms of significance of nestedness and modularity, only the latter are shown. In most of these transformed matrices, the 0 values correspond to an absence (or almost absence) of infection, of symptoms or to a lack of effect on plant health, and the 9 values correspond (or are close) to the maximal possible pathogenicity values.

To test if the matrices could fit with the inverse-matching-allele model (Fig. 1), we also analyzed the “reverse matrices”, where 0 and 9 correspond to the minimal and maximal plant resistance classes, respectively. Methods to estimate nestedness and modularity are detailed in Weitz et al. (2013). Whereas many algorithms can measure the nestedness of matrices containing binary data (0 and 1), only two algorithms were available for matrices containing quantitative numeric data: the weighted nestedness metric based on overlap and decreasing filling (*wNODF* algorithm) (Almeida-Neto et al. 2008) and the weighted-interaction nestedness estimator (*WINE* algorithm) (Galeano et al. 2009). In the R software, the ‘nested’ and ‘wine’ functions were used to estimate the *wNODF* and *WINE* scores, respectively. Because none of the module detection algorithms developed to date provide consistently optimal results in all matrices (Aldecoa and Marín 2013), we used seven different algorithms implemented into the R software for modularity analyses: the *edge betweenness* (Newman and Girvan 2004), *fast greedy* (Clauset et al. 2004), *label prop* (Raghavan et al. 2007), *leading eigenvector* (Newman 2006), *louvain* (Blondel et al. 2008), *spinglass* (Newman and Girvan 2004; Reichardt and Bornholdt 2006; Traag and Bruggeman 2009) and *walktrap* (Pons and Latapy 2006) algorithms (see Supplementary Methods 2 for details).

To help the interpretation of results, we used simulations (which provided test matrices) to compare the performances (type I and type II error rates) of the nestedness and modularity algorithms (Supplementary Methods 2; Supplementary Tables S1 to S18). To determine the statistical significance of the patterns (nestedness or modularity) of the plant-parasite interaction matrices, the actual or test matrices were compared to matrices generated under seven different null models (Supplementary Methods 2). All null-model matrices possess the same dimensions as the actual or test matrix to which they are compared. They differ by the constraints applied for their generation. Under the N null model, matrices are randomly generated ensuring that the total sum of the cells and the number of zero-valued cells are the same as in the actual matrix. Thus, neither the marginal sums of rows or columns, nor the positions of zero-valued cells are constrained. The C1 and R1 null models are generated in the same manner but column by column and row by row, respectively. Under the B null model, matrices with identical column and row marginal sums as the actual matrix are generated using Patefield’s (1981) algorithm. This is the only null model where the number of zero-valued cells can differ from that of the actual matrix. Under the S null model, the cell values are shuffled in the matrix, with no constraints on row or column marginal sums. Finally, under the C2 and R2 null models, the cell values are shuffled column by column and row by row, respectively. We also considered simultaneously the C1 and R1 (or C2 and R2) null models, providing null models CR1 and CR2 in evaluation of test matrices. In summary, these null models comprise the most commonly used in nestedness or modularity analyses and span a large diversity of constraints, N and S being the least constrained, B being the most constrained and the others in between.

Two modularity algorithms (*walktrap* and *label prop*) provided modularity estimates of 0 (or near 0) for almost all actual matrices and associated null models. Moreover, almost all simulations also provided modularity estimates of 0 with these algorithms, hampering the evaluation of type I and type II error rates (Supplementary Methods 2). We could also not evaluate the performance of the *leading eigenvector* algorithm because it did not converge towards a modularity estimate in many simulated matrices. Consequently, these three algorithms were not considered for further analyses.

### Linear model analyses

Using the R software, we analysed two different linear models to evaluate the contribution of the plant-parasite interaction to the pathogenicity trait variance in the matrix data.

Using the mean pathogenicity value for each plant-parasite genotype pair, we analysed the following model: ‘pathogenicity’ ∼ ‘parasite strain’ + ‘plant accession’ that does not include any interaction term. Using independent individual pathogenicity values for each plant-parasite genotype pair, we analysed the following model: ‘pathogenicity’ ∼ ‘parasite strain’ + ‘plant accession’ + ‘parasite strain × plant accession’. The latter model could be applied only to 26 of the 32 matrices. For matrices 1 to 4 with the latter model, the linear model was applied after taking into account a significant effect of the experimenters who performed the measures of the lesion sizes (residues of the linear model ‘pathogenicity’ ∼ ‘experimenter’). For most datasets, the model hypotheses (normality of residues, homoscedasticity) were not satisfied. Consequently, we focused on the model fit (ω^2^, the unbiased estimators of the parts of variance explained by the parasite strain, the plant accession and/or their interaction) rather than on the statistical significance levels or on the comparison of mean pathogenicity values between effect levels.

## Data accessibility

Data are available online: http://doi.org/10.5281/zenodo.5167270

## Supplementary material

Script and codes are available online: http://doi.org/10.5281/zenodo.5167270

Supplementary Methods 1 and 2 (including Supplementary Tables S1 to S18 and Supplementary Figures S1 and S2) and Supplementary Tables S19 to S22 are available online: https://doi.org/10.1101/2021.03.03.433745

## Acknowledgements

Marie-Claire Kerlan and Lionel Renault are acknowledged for there help to produce matrix number 25 and Anne Massire, Ghislaine Nemouchi, Thérèse Phaly, Bruno Savio and Patrick Signoret for their assistance to produce matrix number 17. We thank Amine Slim from the National Gene Bank of Tunisia (NGBT) for providing seeds of the durum wheat landrace “Mahmoudi Joumine” used to build the matrices 12 and 13, and we thank Aurélie Ducasse and Johann Confais for their help in acquiring phenotypic data on the wheat-*Zymoseptoria tritici* pathosystem found in matrices 14 and 15. We thank Isabelle Demeaux (INRAE, SAVE) for providing technical assistance with the downy mildew/grapevine pathosystem. Anne Quillévéré-Hamard, Gwenola Le Roy and Christophe Le May are acknowledged for having co-supervised, managed and/or significantly contributed to the production of matrices 19 and 20. We thank Loup Rimbaud and Emmanuel Szadkowski (INRAE, PACA) for their comments on an earlier version of the manuscript and Michel Pitrat (INRAE, PACA) for his help for analyses of matrices 10 and 11. We thank the staff of the INRAE CRB-Leg (https://www6.paca.inrae.fr/gafl/CRB-Legumes) who maintained the pepper and melon germplasm collections of the GAFL research unit, and of the INRAE experimental facilities of the Plant Pathology research unit (https://doi.org/10.15454/8DGF-QF70), the GAFL experimental unit and the PHENOTIC core facility in Angers (https://doi.org/10.15454/U2BWFJ) who ensured the production of the plants and maintenance of plant-growth facilities that allowed us to do this work. We thank the staff of the INRAE experimental facilities of IGEPP for having provided and managed equipment for the experiments.

The research was supported by the French National Research Agency (ANR) programs BIOADAPT (grant no. ANR-12-ADAP-0009-04), ArchiV (grant no. ANR-18-CE32-0004-01), CEDRE (grant no. ANR-05-PADD-05) and PeaMUST (grant no. ANR-11-BTBR-0002), the PROGRAILIVE project (grant RBRE160116CR0530019) funded by the Bretagne region, France and European FEADER grants, the fundings of the Institut Carnot PLANT2PRO and the Comité Interprofessionnel des Vins de Bordeaux (CIVB), the INRAE departments “Santé des Plantes et Environnement” (project APÔGÉ and PhD thesis of Safa Ben Krima) and “Génétique et Amélioration des Plantes”, the INRAE métaprogramme SMaCH (Sustainable Management of Crop Health), the French Ministry of Agriculture and Food for projects “Recherche et mise au point de méthodes pour évaluer des résistances variétales durables à des agents pathogènes” (CTPS project C2008-29), “Nouvelles sources de résistance à *Aphis gossypii* chez le melon” (CTPS project C06/02) and “Caractérisation de la virulence de *Podosphaera xanthii*, agent causal de l’oïdium du melon, et développement d’un système de codification des races” (CTPS project C-2012-10). UMR1290 BIOGER benefits from the support of Saclay Plant Sciences-SPS (ANR-17-EUR-0007).

Version 4 of this preprint has been peer-reviewed and recommended by *Peer Community In Evolutionary Biology* (https://doi.org/10.24072/pci.evolbiol.100132).

