## Supplementary material for "The quasi-universality of nestedness in the structure of quantitative plant-parasite interactions": Suppl Methods 1 and 2; Suppl. Tables 19 to 22: Suppl_Methods_1.docx

**Supplementary Methods 1: Characteristics of experimental datasets and phenotyping procedures**

Matrices 1 to 4: *Pseudomonas syringae-Prunus armeniaca* (apricot)

Nine strains of *Pseudomonas syringae*, the causal agent of bacterial canker of apricot, were inoculated on dormant tissues of twenty apricot cultivars chosen according to their differential susceptibility in orchard conditions. The strains were chosen mainly within phylogroups 1 and 2, the most abundant groups of *P. syringae* in contaminated apricot orchards in France (Parisi *et al*. 2019). Seven strains were isolated from symptomatic trees and two in crop debris and soil. Bacterial inoculum was prepared by cultivation on King's B medium for 48h at 24°C. The concentration of the bacterial suspension was adjusted at 10^8^ CFU.ml^-1^. A volume of 25 μl of inoculum was deposited at the level of a wound made superficially with a scalpel on the bark of one-year-old twigs grown in orchard. Five months after inoculation, twigs were removed and the length of flat zone around the inoculation point at the surface of the shoot (matrices 1 and 2) and the length of browning zone around the inoculation point below the bark of the shoot (matrices 3 and 4) were measured. Two independent tests were performed in 2017 (matrices 1 and 3) and 2018 (matrices 2 and 4).

Matrix 5: *Puccinia hordei-Hordeum vulgare* (barley)

Fourteen *Puccinia hordei* isolates (from Europe, Morocco, Israel and the USA) were inoculated on a differential series of 12 *H. vulgare* lines carrying different *Rphq* QTLs (González et al. 2012). The first seedling leaves of each barley line were inoculated with ≈240 spores/cm^2^. The relative latency period (RLP) (Table 3 in González et al. 2012) was estimated by the number of hours from inoculation to the moment at which 50% of the ultimate number of uredinia was visible.

Matrices 6 to 8: *Venturia inaequalis-Malus domestica* (apple tree)

Grafted plants of different apple tree accessions (*Malus domestica*) carrying resistance QTLs (*T1*, *F11*, *F17*, *F11* + *F17* or *T1* + *F11* + *F17*) or no resistance QTL were inoculated in controlled conditions with isolates of *Venturia inaequalis*, a fungal pathogen responsable of apple scab. The percentage of sporulating leaf area was assessed from 8 to 21 days post inoculation (dpi) on a scale with eight levels: 0 = no visible symptom, 0.5 = 0–1%, 3 = 1–5%, 7.5 = 5–10%, 17.5 = 10–25%, 37.5 = 25–50%, 62.5 = 50–75%, and 87.5 = 75–100%.

Matrix 6 (Laloi et al. 2017) consisted of interactions between 10 *V. inaequalis* isolates sampled in one orchard (Angers, France) on apple trees carrying *T1*, *F11*+*F17*, *T1*+*F11*+*F17* or no QTL and 14 apple tree accessions carrying the matching resistance QTL or no QTL.

Matrix 7 (Caffier et al. 2016) consisted of interactions between 14 *V. inaequalis* isolates sampled in one orchard (Angers, France) on apple trees carrying or not *T1* and 12 apple tree accessions carrying or not *T1* (with six accessions for each of the two classes). Matrices 6 and 7 represent the Area Under the Disease Progress Curve (AUDPC) of the percentage of sporulating leaf area from eight to 21 dpi.

Matrix 8 (Caffier et al. 2014) consisted of interactions between 24 *V. inaequalis* isolates sampled in two orchards (Lanxade and Villeneuve d’Ascq, France) on apple trees carrying *F11*, *F17*, *F11*+*F17* or no QTL and eight apple tree accessions carrying the matching QTL or no QTL (with two accessions for each of the four classes). Matrix 8 represents the percentage of sporulating leaf area 14 dpi.

Matrix 9: *Botrytis cinerea-Solanum lycopersicum* / *Solanum pimpinellifolium* (tomato)

Leaves of 12 tomato accessions (six domesticated accessions of *Solanum lycopersicum* and six accessions of the close wild relative *S. pimpinellifolium*) were infected with single droplets of spore suspensions of 94 *B. cinerea* isolates. The size of lesions was measured from digital images 72 hours after inoculation (Soltis et al. 2019). One isolate was poorly infectious on all tomato accessions (grade 0 after data transformation) and was withdrawn.

Matrices 10 and 11: *Podosphaera xanthii-Cucumis melo* (melon)

Nineteen melon lines were inoculated with 26 *Podosphaera xanthii* isolates collected in 2013 and 2014 in melon, squash, watermelon and cucumber crops in Southern Europe or Northern Africa (France, Spain, Italy, Morocco, Turkey, Greece). Each *P. xanthii* isolate was propagated on cotyledons of *Lagenaria* *ciceraria* for seven days and spores were blown on eight leaf disks per melon line-*P. xanthii* isolate combination using an inoculation tower (Perchepied et al. 2005). Sporulation intensity was scored 14 days after inoculation and data were transformed in percentage of leaf disk surface using the class mean as suggested by Nicot et al. (2002): 0 = 0%, 1 = 2.5%, 2 = 7.5%, 3 = 17.5%, 4 = 37.5%, 5 = 67.5%, 6 = 82.5%, 7 = 92.5%, 8 = 97.5%, and 9 = 100%. The mean score for melon accession - *P. xanthii* isolate combinations was reported in matrix 10. For matrix 11, 31 isolates were inoculated to 19 differential lines on leaves of entire plants. The sporulation intensity was scored similarly as for leaf disks using a 0 to 9 scale. The mean score for melon accession - *P. xanthii* isolate combinations was reported in matrix 11.

Matrices 12 to 15: *Zymoseptoria tritici-Triticum aestivum* (bread wheat) or *T. turgidum* subsp*. durum* (durum wheat)

Matrices 12 and 13 were built by inoculating 12 lineages from a durum wheat landrace called Mahmoudi with 15 *Zymoseptoria tritici* isolates. The 12 plant lineages were fixed from individivuals coming from a single field at Joumine in Tunisia and corresponded to 12 different multilocus genotypes (MLGs) as defined previously by Ben Krima et al. (2020). The 15 isolates were collected *in situ* either from the landrace Mahmoudi or from the cultivar Karim, at Joumine in 2018. Matrices 14 and 15 were built by inoculating 15 bread wheat cultivars (*Triticum aestivum*), 12 of which carrying different *Stb* resistance genes, with 98 *Z. tritici* isolates collected mostly on cultivars Apache and Premio, all over France between 2009 and 2010. These bread wheat cultivars belong to a series of differential genotypes used to characterize the pathogenicity of *Z. tritici* isolates. All wheat-*Z. tritici* pairwise confrontations were evaluated under controlled conditions, in growth chambers at 18°C/22°C night/day and 16 hours light at 300 µmol.m^-2^.s^-1^. The first true leaf of 16-day-old seedlings were marked with a black felt to delimit a 7.5 cm length that was inoculated with a solution of water containing 10^6^ spores.mL^-1^ and one drop of Tween®20 per 15 mL. The inoculum was applied with a square-tipped flat paintbrush six times on each leaf, repeated twice. After inoculation the plants were placed in transparent polyethylene bags for 72 hours to initiate infection. At 10 dpi, *i.e.* before the appearance of symptoms, leaves above the inoculated leaf were cut to homogenize light exposure. Visual estimations of necrotic leaf area and sporulating leaf area were done at 14 dpi, 20 dpi and 26 dpi for matrices 12 and 13, and only once at 21 dpi for matrices 14 and 15. For matrices 12 and 13, these observations were used to calculate, for each plant lineage-isolate combination, an area under disease progress curve (AUDPC) for the percentages of necrotic and sporulating leaf areas. The interactions for matrices 12 and 13 were evaluated on three leaves repeated twice in time (total of six leaves) and for matrices 14 and 15 on three leaves repeated thrice in time (total of nine leaves).

Matrices 16, 17 and 17b: *Phytophthora capsici-Capsicum annuum* (pepper)

To build matrix 16, the pathogenicity of six isolates of *Phytophthora capsici*, the causal agent of root and crown rot of chilli and bell peppers, collected in pepper fields in Algeria was measured in ten *Capsicum annuum* cultivars (F_1_ hybrids or inbred lines) (Messaouda et al. 2015). Six plants per accession were inoculated by depositing a plug of 4 mm in diameter of mycelium of *P. capsici* cultivated on V8 medium on the fresh section of the primary stem extemporaneously decapitated (Lefebvre and Palloix 1996). Inoculated plants were kept in a growth chamber under controlled conditions with 12h photoperiod, a temperature of 22 ± 2°C and 100% relative humidity. *P. capsici* progresses to the bottom of the stem causing a necrosis of the stem. The length of stem necrosis at 15 dpi is reported in matrix 16.

For matrix 17, 53 accessions of *C. annuum* were inoculated by six isolates of *P. capsici*. The *C. annuum* accessions originated from 20 countries from America, Europe, Asia and Africa, and included accessions that had different levels of partial resistance to isolate *P. capsici* ‘Pc101’ and a few susceptible accessions. The six *P. capsici* isolates were isolated from pepper plants in France and Turkey, were of A1 mating type and differed in pathogenicity. A minimum of six plants per accession, seven-eight week old, were inoculated as described for matrix 16. Inoculated plants were kept in a growth chamber under controlled conditions with a photoperiod of 12h at 24°C under artificial light and 22°C at obscurity. The length of stem necrosis was measured six times from three to 21 dpi and the Area Under the Disease Progress Curve (AUDPC) of necrosis length was considered in matrix 17.

Because matrix 17 contained a large number of zero-values cells, matrix 17b was derived by withdrawing redundant columns and columns entirely made of zero-valued cells.

Matrix 18: *Phytophthora infestans-Solanum lycopersicum*

Matrix 18 was built by inoculating eight *Solanum* sp. accessions with seven isolates of *Phytophthora infestans*, the causal agent of tomato late blight. The accessions consisted of three inbred lines of cultivated tomato (*Solanum lycopersicum*) and five accessions of the wild relative species *S. pimpinellifolium*, *S. habrochaites* and *S. pennellii*. Some of them are known to carry the *Ph-1*, *Ph-2* or *Ph-3* genes, controlling resistance to races 0, 1 and 2 of *P. infestans*, respectively. The *P. infestans* isolates were collected on tomato or potato plants in France and Poland and were chosen because they varied in mating type (A1 or A2) and differed in pathogenicity. Mycelium was grown on pea juice-based agar medium for 10 days and six plants per accession, 3-4 week old, were inoculated using the protocol described for matrix 16 (Danan et al. 2009). Inoculated plants were kept in a growth chamber under controlled conditions with a photoperiod of 14h at 21°C under artificial light and 17°C at obscurity. High humidity was maintained by artificial mist. Stem necrosis length was scored four times from three to 14 dpi and the AUDPC was calculated.

Matrix 19: *Aphanomyces euteiches*-Fabaceae (pea, vetch, faba bean, alfalfa)

Eight accessions from four leguminous species (pea, alfalfa, vetch, faba bean), which previously showed various levels of resistance, were inoculated with 34 *Aphanomyces euteiches* isolates sampled from the main French pea growing regions in a growth chamber (thermo period: 25/23°C and 16h photoperiod). Seven-day-old plants (5 plants * 4 replicates * 2 experiments for each accession-isolate combination) were inoculated by applying 5 mL of a zoospore suspension adjusted to 5.10^3^ spores / mL. After inoculation, the vermiculite substrate was saturated with water to provide favorable conditions for infection. After 10 days, the plants were carefully removed from the vermiculite, the roots were washed in tap water and disease severity (DS) was scored on each plant using a 0–5 scale: 0 = no symptoms; 1 = traces of discoloration on the roots (<25%); 2 = discoloration of 25 to 50% of the roots; 3 = discoloration of 50 to 75% of the roots; 4 = discoloration of >75% of the roots; 5 = dead plant. ANOVA was performed with the DS score as the dependent variable, the *A. euteiches* isolate and the plant accession as fixed factors and the replicate and experiment as random factors. From the ANOVA, least square means (LSmeans) were calculated for each *A. euteiches* isolate-plant accession combination. In the present study, LSmeans values of root DS scores were analysed. More details are provided in Quillévéré-Hamard et al. (2018).

Matrix 20: *Aphanomyces euteiches*-*Pisum sativum*

Ten pea accessions were inoculated with 43 *A. euteiches* isolates sampled from the main French pea growing regions in a growth chamber. The ten pea accessions consisted of (i) eight Near-Isogenic-Lines (NILs) carrying one, two, three or five resistance alleles at main QTLs, in a common genetic background and (ii) two control lines, including one susceptible variety and one highly resistant line. The experimental design, inoculation procedure, disease scoring scale and statistical analysis were similar to that described for matrix 19, except for inoculum concentration (2.10^2^ spores / mL) and the scoring date (seven days after inoculation). LSMean values of root DS scores were used. More details are provided in Quillévéré-Hamard et al. (2021).

Matrices 21 and 22: *Plasmopara viticola-Vitis vinifera* (grapevine)

A set of 33 *Plasmopara viticola* strains, the causal oomycete of grapevine downy mildew, was inoculated on eight grapevine varieties. The host panel was constituted of seven grapevine varieties carrying the main resistance factors currently used in European breeding programs (*Rpv1*, *Rpv3.1*, *Rpv3.2*, *Rpv5*, *Rpv6*, *Rpv10* and *Rpv12*) and one susceptible variety (Chardonnay). Cuttings from these varieties were grown in a glasshouse under natural photoperiod. Each strain-variety combination was replicated on five leaf discs from five different plants that were excised in the fourth leaf below the apex. Leaf discs were sprayed with 4 mL of a suspension of 10^5^ / mL sporangia of *P. viticola*. They were incubated in a climatic chamber for six days at 18°C with 12h/12h light/dark photoperiod. At six dpi, necrosis was rated on a scale of 0 to 4, based on the number of necroses counted per leaf disk (0 = no necrosis; 1 = <10 necroses; 2= from 10 to 30 necroses; 3= from 30 to 60 necroses; 4= > 60 necroses) (matrix 21) and sporulation was assessed on leaf discs by automatic image analysis (number of white pixels on the total leaf disc area) (matrix 22).

Matrices 23 and 24: *Aphis gossypii-Cucumis melo*

Matrices 23 and 24 were obtained through assessment of the resistance of 13 melon accessions to nine aphid (*Aphis gossypii*) clones (Boissot et al. 2016). The host panel consisted in twelve partially-resistant lines originating from Africa, India, China, Asia and Far East Asia, Mediterranean basin and North America and a susceptible cultivar originating from Mediterranean basin. Two lines were wild accessions and the others from breeding programs. They contained at least one to three homologs of *Vat*, a gene conferring resistance to *A. gossypii*. The 13 melon lines belonged to three genetic groups representative of melon diversity (Boissot et al., submitted). The aphid panel consisted in nine clones collected in France and French West Indies. Except clone NM1 that was observed on plant species belonging to six families, the clones have been observed exclusively (or almost exclusively) on cucurbit plants and belong to the same genetic cluster.

For phenotyping, ten adult aphids were deposited on melon plantlets. Three days later, the number of aphids remaining on the plantlets was recorded as the ‘Acceptance’ parameter (matrix 23). Seven days after aphid deposition, the adults were counted, and the density of nymphs was estimated on a scale of 0 to 6. The ‘Colonization’ parameter was calculated as [density of nymphs + ln(number of adults + 0.001)] (matrix 24). The ‘Acceptance’ and ‘Colonization’ parameters were collected for at least eight plantlets of each melon accession. Each test was conducted with one aphid clone on a subset of melon accessions.

Matrices 25 and 26: *Globodera pallida-Solanum tuberosum* (potato)

Matrix 25 was obtained through the inoculation of 20 populations of the potato cyst nematode *Globodera pallida* on ten potato accessions. Those potato accessions were characterized by different levels of quantitative resistance. A susceptible potato cultivar, Désirée, was also used as a control. Among the 20 *G. pallida* populations, 14 came from South-America (Peru and Chile) and six from Europe. To perform *G. pallida* inoculation, ten cysts were locked in a tulle bag and placed in a pot three-quarter filled with a soil mixture free of cysts (2/3 sand and 1/3 natural field soil). Four replicates were performed for each potato accession - *G. pallida* population combination, *i.e.* for each *G. pallida* population, four bags were inoculated to four tubers of the same potato accession. One potato tuber was planted per pot and covered with the same soil mixture. Potato plants grew in the greenhouse, under controlled conditions (15°C night during 8h and 20°C day during 16h), for 120 days. After 120 days, newly formed cysts were extracted from the soil, using a Kort elutriator. The number of newly formed cysts was counted using a magnifying stereomicroscope, and divided by the number of newly formed cysts produced on the susceptible cultivar Désirée (relative value).

For matrix 26, the measured fitness trait was the hatching of cysts which is induced by host root exudates. It was produced using a cross-hatching assay between 13 populations of *G. pallida* and root exudates from 12 wild potato accessions, belonging to species *Solanum huancabambense*, *S. mochiquense*, *S. sogarandinum*, *S. ambosinum*, *S. medians*, *S. pampasense*, *S. santalallae*, *S. marinasense*, *S. sparsipilum*, *S. raphanifolium*, *S. limbaniense* and *S. leptophyes*, to test the hypothesis of local adaptation between Peruvian *G. pallida* populations and Peruvian wild potato accessions (Gautier et al. 2020). All details about *G. pallida* populations, root-exudates and the *in vitro* hatching assay are available in Gautier et al. (2020). Briefly, three cysts of each population were put on a sieve in 1.5 mL of root exudates (with four to five replicates) and after 30 days, the number of hatched juveniles was counted. At the end of the experiment, cysts were crushed and the number of unhatched viable eggs was counted, in order to calculate a hatching percentage.

Matrices 27 to 32: *Potato virus Y-Capsicum annuum*

The *Capsicum annuum* accessions were doubled-haploid lines issued from the F_1_ hybrid between accessions Perennial, carrying several *Potato virus Y* (PVY) resistance QTLs, and the susceptible accession Yolo Wonder. They were chosen based on the lack of a major-effect resistance gene but contrasted levels of quantitative resistance (Caranta et al. 1997). The PVY populations were issued from cDNA clones of isolates SON41p and LYE84.2 and recombinants between these two cDNA clones (Montarry et al. 2012). *Capsicum annuum* accessions were mechanically inoculated with the different PVY populations and the virus load at the systemic level was estimated one month post inoculation by quantitative DAS-ELISA as described in Quenouille et al. (2014) (matrices 27 and 28). In addition, the area under the disease progress curve (AUDPC) was calculated using a semi-quantitative scoring scale as in Caranta et al. (1997) (matrices 29 and 30) and the dry weight of infected relative to mock-inoculated plants was estimated as in Montarry et al. (2012) (matrices 31 and 32). Matrices 27, 29 and 31 on the one hand and matrices 28, 30 and 32 on the other hand correspond to two independent experiments with slightly different sets of PVY populations.
