## Supplementary material for "The quasi-universality of nestedness in the structure of quantitative plant-parasite interactions": Suppl Methods 1 and 2; Suppl. Tables 19 to 22: Suppl_Methods_2.docx

**Supplementary Methods 2: Comparison of the performance of algorithms used to detect nested or modular matrix structures**

We used simulations to evaluate the type I error rates and the statistical power (*i.e.* 1-*E*_2_, where *E*_2_ is the type II error rate) associated to two algorithms used to detect nestedness and seven algorithms used to detect modularity.

**1. Simulations**

1.1. Nestedness algorithms

The *wNODF* (weighted nestedness metric based on overlap and decreasing filling) estimator (Almeida-Neto and Ulrich 2011) is a weighted version of the nestedness measure originally proposed by Almeida-Neto et al. (2008). Values of 0 indicate non-nestedness, those of 100 perfect nesting (for example Fig. 3A). *wNODF* quantifies whether a given sequence of columns or rows shows a gradient of decreasing marginal totals.

The weighted-interaction nestedness estimator (*WINE* algorithm) (Galeano et al. 2009) is based on the concept of estimating nestedness through the calculation of a weighted Manhattan distance from each of the matrix cells containing a value >0 to the cell corresponding to the intersection of the row and column with the lowest marginal totals. The statistical significance of this nestedness index value is tested against a null model that constrains matrix fill to observed values, retains the distribution of number of events in the links but does not constrain marginal totals. The *WINE* index varies usually from 0 (low nestedness) to 1 (high nestedness), though values higher than 1 can sometimes be obtained.

*WINE* overcomes the limitations of some previous indices, being independent from matrix size and fill (i.e. frequency of non-zero-valued cells) (Galeano et al. 2009).

1.2. Modularity algorithms

A number of algorithms allows to detect modules in a network, i.e. groups of densely connected nodes with fewer connections across groups. For an increasing number of modules, these methods estimate a maximum modularity score which varies from 0 (non-modular, i.e. random matrix) to 1 (high modularity). The highest modularity score across all module numbers allows to define both the optimal number of modules and the distribution of plant and parasite genotypes in these modules.

We used the *edge betweenness* (Newman and Girvan 2004), *fast greedy* (Clauset et al. 2004), *label prop* (Raghavan et al. 2007), *leading eigenvector* (Newman 2006), *louvain* (Blondel et al. 2008), *spinglass* (Newman and Girvan 2004; Reichardt and Bornholdt 2006; Traag and Bruggeman 2009) and *walktrap* (Pons and Latapy 2006) algorithms. We refer to the original articles for a presentation of the methods and to Weitz et al. (2013) for an overview. Four of these seven algorithms could be applied to our actual matrices and their performances evaluated by our simulation approach. We could neither analyze the modularity of our actual matrices with the *walktrap* and *label prop* algorithms nor evaluate the performances of these algorithms because the modularity scores of almost all test matrices and null-model matrices were equal to zero. We could also not evaluate the performance of the *leading eigenvector* algorithm because it did not converge towards a modularity estimate for many simulated matrices.

1.3. Overall simulation procedure

For each evaluation of type I or type II errors, 100 matrices were simulated (see description of these "test matrices" in section 1.4), either with expected nested or modular structures (for type II errors) or with no nested or modular structures expected (for type I errors). Each of these test matrices was compared with 100 random matrices simulated under various null models (see description of these "null-model matrices" in section 1.5) for their degree of nestedness (or modularity) to calculate one probability value (p-value) corresponding to the significance of nestedness (or modularity) of the test matrix. A test matrix is significantly nested (or modular, respectively) when its degree of nestedness (or modularity) exceeds that of ‘a majority’ of the null-model matrices. The frequency of p-values <0.05 among the 100 test matrices (F_0.05_) provided an estimation of type I errors for nestedness/modularity. In other words, a 10% level of type I errors for nestedness significance assessment means that 10 of 100 test matrices simulated with no expected nested structure had a p-value strictly below 0.05, i.e. the nestedness estimated for each of these 10 test matrices was strictly higher than the nestedness estimated for 95% or more of 100 simulated null-model matrices used for comparison.

Similarly, an estimation of the statistical power of nestedness/modularity estimation methods is provided by F_0.05_ obtained with the appropriate test matrices, i.e. (1-F_0.05_) corresponds to the level of type II errors.

This approach also provides an evaluation of type I errors for the detection of anti-nestedness (or anti-modularity). A test matrix is significantly anti-nested (or anti-modular, respectively) when its degree of nestedness (or modularity) is lower than that of ‘a majority’ of the null-model matrices. The frequency of p-values > 0.95 among the 100 test matrices (F_0.95_) provided estimations of type I errors for anti-nestedness/anti-modularity. In other words, a 10% level of type I errors for anti-nestedness significance assessment means that 10 of 100 test matrices simulated with no expected nested (or anti-nested) structure had a p-value strictly above 0.95, i.e. the nestedness estimated for each of these 10 test matrices was strictly higher than the nestedness estimated for 95% or more of 100 simulated null-model matrices used for comparison.

We did not estimate the statistical power of anti-nestedness/anti-modularity estimation methods because we did not simulate test matrices with *a priori* anti-nested or anti-modular structures.

1.4. Test matrices

For all simulations, we used 10 × 10 matrices. These dimensions are close to the median of dimensions of the actual plant-parasite matrices analysed.

To evaluate the statistical power of nestedness methods, five patterns (M1 to M5; Fig. S1) were used to simulate test matrices. Similarly to the actual matrices, these test matrices contained integer values from 0 to 9 (except M1, used for comparison with M2, that contained integer values from 0 to 5) chosen at random but in different intervals for the different cells in order to obtained matrices with expected nested patterns.

To evaluate the statistical power of modularity methods, six patterns (M6 to M11; Fig. S2) were used to simulate test matrices. These test matrices contained integer values from 0 to 9 chosen at random but in different intervals for the different cells in order to obtained matrices with expected modular patterns.

To evaluate the type I errors of nestedness and modularity methods, we simulated test matrices with no expected nested or modular patterns. We started by simulating test matrices following patterns M1 to M5. Then, for each obtained matrix, the cell values were either randomized globally in a matrix of same dimensions (providing the corresponding patterns M1S to M5S) or the cell values in each row of the matrix were randomly permuted (providing the corresponding patterns M1R to M5R). Due to symmetry reasons, similar results are obtained by permuting the cell values in the matrix columns.

1.5. Null-model matrices

To evaluate the statistical significance of the nestedness (or modularity) degrees of each test matrix, 100 matrices were simulated under seven different null models, and their nestedness (or modularity) degrees were estimated for comparison. All null-model matrices had the same dimensions as the test matrices (10 × 10). The different null models had different types of constraints and were generated from the actual matrix using the ‘permatfull’ function in the ‘bipartite’ R package.

Under the N null model (permatfull options fixedmar = "none" and shuffle = "both"), new matrices are randomly generated ensuring that the total sum of the cells and the number of zero-valued cells are the same as in the actual matrix. Thus, neither the marginal sums of rows or columns, nor the position of zero-valued cells are constrained.

The C1 and R1 null models (permatfull options fixedmar = "columns", shuffle = "both" and fixedmar = "rows", shuffle = "both", respectively) are generated in the same manner but column by column and row by row, respectively.

To reduce the type I errors, we defined the probability value of the CR1 model as the highest (*i.e.* least significant) among those obtained under the C1 and R1 null models for a given test matrix.

Under the B null model (permatfull options fixedmar = "both" and shuffle = "both"), a matrix with identical column and row marginal sums as the actual matrix is generated using Patefield's (1981) algorithm. This is the only null model where the number of zero-valued cells can differ from that of the actual matrix.

Under the S null model (permatfull options fixedmar = "none" and shuffle = "samp"), the cell values are shuffled in the matrix, with no constraints on row or column marginal sums.

Under the C2 and R2 null models (permatfull options fixedmar = "columns", shuffle = "samp" and fixedmar = "rows", shuffle = "samp", respectively), the cell values are shuffled column by column and row by row, respectively.

For a given test matrix, the maximal p-value obtained with either C2 or R2 was also considered (null model CR2).

1.6. Implementation

All simulations and statistical analyses were performed using the R program version 3.5.1. The ‘bipartite’ and ‘igraph’ packages were used for nestedness and modularity analyses, respectively. The ‘permatfull’ function was used to generate the null-model matrices B, N, C1 and R1. The ‘wine’ and ‘nested’ functions were used to estimate matrix nestedness with the *WINE* and *wNODF* algorithms, respectively. The ‘cluster_edge_betweenness’, ‘cluster_fast_greedy’, ‘cluster_label_prop’, ‘cluster_leading_eigen’, ‘cluster_louvain’, ‘cluster_spinglass’ and ‘cluster_walktrap’ functions were used to estimate matrix modularity with the corresponding algorithms.

**2. Results**

2.1. Type I errors of nestedness

With test matrix patterns M1S to M5S, where cell values were shuffled in the matrix, relatively few type I errors (<10%) were obtained with both methods (*WINE* and *wNODF*) and with all null models, with a few exceptions for *wNODF* with the B null model (Supplementary Tables S1 and S2). In contrast, with test matrix patterns M1R to M5R, where cell values were randomized in rows but show a value gradient in the columns, many false positive nestedness were evaluated, especially with null models N, C1, S and C2. The R1, R2, CR1 and CR2 null models provided satisfactory results with <10% false positives in almost all cases. For symmetry reasons, if the cell values of the test matrices had been randomized in columns (instead of rows) and showed a value gradient in the rows, the C1, C2, CR1 and CR2 null models would have been satisfactory, but not the R1 and R2 null models (data not shown). Consequently, to avoid false positives in cases of cell value gradients either in rows or columns, the best null models are CR1 and CR2, with 2 to 3% false positives on average for all test matrices at the 0.05 significance threshold.

Comparison between matrix patterns M1S and M2S (or between M1R and M2R), which share a similar structure but where cell values vary from 0 to 5 and 0 to 9, respectively, allows estimating the impact of the scale of values used to transform the initial matrices. The type I error rates of M1S and M2S were almost identical, whereas it was higher for M2R than for M1R. However, similar type I error rates were observed for M2R and M1R with the best null models (CR1 and CR2).

2.2. Type I errors of anti-nestedness

Whatever the test matrix, the null model and the algorithm, few type I errors (<10%) were obtained for anti-nestedness in almost all cases, except for the *WINE* method with null model B (Supplementary Tables S3 and S4). M1S and M2R from one side, and M1R and M2R from the other side provided similar type I error rates.

2.3. Statistical power of nestedness

Except for the B and CR1 null models, the statistical power of both nestedness algorithms was >30% in most cases (Supplementary Tables S5 and S6). The CR2 null model had the highest power varying from 38 to 100%, depending on the algorithm and on the nestedness degree of the test matrix. The power was usually higher with the *WINE* than with the *wNODF* algorithm when compared for the same test matrix and the same null model.

M2 had a higher power than M1 for null models with the highest power (N, S and especially CR2). It was the opposite for models with the lowest power (B and CR1).

2.4. Type I errors of modularity

For the *edge betweenness*, *fast greedy* and *louvain* algorithms, only the null models S, C2, R2 and CR2 provided low type I errors (≤0.10) with all test matrices (Supplementary Tables S7 to S9). For the *spinglass* algorithm, all null models were highly satisfactory with all test matrices (≤0.01 type I errors for modularity significance; Supplementary Table S10).

M1S vs M2S and M1R vs M2R: much more false positives were observed with model pattern M2S (or M2R) than with M1S (or M1R). However, similar performances were observed between these model patterns for the best null models (S, C2, R2 and CR2) and for the best overall method (*spinglass* algorithm), whatever the null model.

2.5. Type I errors of anti-modularity

For the *edge betweenness* algorithm, all null models except S and C2 were highly satisfactory with all test matrices (≤0.1 type I errors for anti-modularity significance in almost all cases; Supplementary Table S11).

For the *fast greedy* and *louvain* algorithms, only the R2 and CR2 null models provided satisfactory results with <10% false positive anti-modularity for all test matrices (Supplementary Table S12 and S13). However, the R2 null model performs similarly to C2 for test matrices that show value gradients in the rows (data not shown) and therefore only the CR2 null model is satisfactory for test matrices with any kind of value gradient (either in rows or columns).

For the *spinglass* algorithm, all null models were highly satisfactory with all test matrices (≤0.01 type I errors for anti-modularity significance; Supplementary Table S14).

M1S and M2R from one side, and M1R and M2R from the other side provided similar type I error rates.

2.6. Statistical power of modularity

In general, whatever the test matrix and the algorithm, the statistical power was higher with null model B, followed by null models N, C1, R1 and CR1 and null models S, C2, R2 and CR2 showed a lower power (Supplementary Tables S15 to S18). The only exception to this tendency is the *fast greedy* algorithm with the M6 test matrix (Supplementary Table S16). The power varied greatly among the test matrices. Whereas the *edge betweenness* algorithm had a quite satisfactory power (>30%) with all test matrices compared to null models B, N, C1, R1 or CR1 (Supplementary Table S15), the *fast greedy* and *louvain* algorithms were efficient to detect significant modularity only with the test matrices showing relatively high modularity degrees (M8 to M11) (Supplementary Tables S16 and S17) and the *spinglass* algorithm was only efficient to detect significant modularity with the test matrix of highest modularity degree (M11) (Supplementary Table S18).

**Supplementary Table S1.** Type I errors of the *WINE* method for nestedness estimation.

| **Nestedness type I error, *WINE* method** | | | | | | | | | |
| --- | --- | --- | --- | --- | --- | --- | --- | --- | --- |
| **Test matrix** | **Null model** | | | | | | | | |
|  | **B** | **N** | **C1** | **R1** | **CR1** | **S** | **C2** | **R2** | **CR2** |
| **M1R** | **0.04***^a^* | 0.11 | 0.24 | **0.05** | **0.02** | 0.18 | 0.27 | **0.06** | **0.01** |
| **M2R** | **0** | 0.23 | **0.35** | **0.02** | **0** | 0.26 | **0.33** | **0.04** | **0.03** |
| **M3R** | **0** | **0.48** | **0.62** | **0.05** | **0.03** | **0.47** | **0.68** | **0.06** | **0.06** |
| **M4R** | **0.45** | 0.25 | **0.49** | **0.09** | **0.09** | **0.37** | **0.55** | **0.06** | **0.04** |
| **M5R** | 0.12 | **0.98** | **1** | **0.09** | **0.09** | **0.99** | **1** | **0.07** | **0.07** |
| **M1S** | **0.01** | **0.03** | **0.05** | **0.04** | **0** | **0.04** | **0.04** | **0.05** | **0** |
| **M2S** | **0** | **0.06** | **0.07** | **0.03** | **0** | **0.07** | **0.06** | **0.08** | **0.02** |
| **M3S** | **0** | **0.04** | **0.03** | **0.04** | **0** | **0.03** | **0.01** | **0.04** | **0** |
| **M4S** | 0.25 | **0.03** | **0.05** | **0.07** | **0.02** | **0.06** | **0.04** | **0.05** | **0.01** |
| **M5S** | **0** | **0.04** | **0.03** | **0.04** | **0.01** | **0.02** | **0.05** | **0.04** | **0** |

*^a^* For each test matrix pattern (table row) and each null model pattern (table column), 100 test matrices were simulated, each of which being compared to 100 null-model matrices. This comparison provided a nestedness significance (p-value) for each simulated test matrix. The frequencies of nestedness p-values < 0.05 (F_0.05_) among the 100 simulated test matrices are indicated in the table. Favorable situations showing relatively few (F_0.05_ ≤ 0.10) false positive nestedness estimations are in bold on grey cells and unfavorable situations showing relatively numerous (F_0.05_ ≥ 0.30) false positive nestedness estimations are in white on black cells.

**Supplementary Table S2.** Type I errors of the *wNODF* method for nestedness estimation.

| **Nestedness type I error, *wNODF* method** | | | | | | | | | |
| --- | --- | --- | --- | --- | --- | --- | --- | --- | --- |
| **Test matrix** | **Null model** | | | | | | | | |
|  | **B** | **N** | **C1** | **R1** | **CR1** | **S** | **C2** | **R2** | **CR2** |
| **M1R** | **0.03***^a^* | 0.15 | 0.23 | **0.06** | **0.03** | 0.12 | 0.24 | **0.07** | **0** |
| **M2R** | **0.04** | 0.22 | **0.36** | **0.06** | **0.02** | 0.18 | 0.29 | **0.01** | **0.01** |
| **M3R** | **0.44** | **0.40** | **0.66** | **0.05** | **0.04** | **0.39** | **0.53** | **0.05** | **0.04** |
| **M4R** | NA*^b^* | NA | NA | NA | NA | NA | NA | NA | NA |
| **M5R** | **0.45** | **0.78** | **0.89** | 0.13 | 0.12 | **0.61** | **0.76** | **0.07** | **0.06** |
| **M1S** | **0.02** | **0.05** | **0.07** | **0.06** | **0.01** | **0.05** | **0.05** | **0.07** | **0.01** |
| **M2S** | **0.04** | **0.07** | **0.08** | **0.08** | **0** | **0.07** | **0.05** | **0.06** | **0.02** |
| **M3S** | **0.85** | **0.06** | **0.06** | **0.08** | **0.01** | **0.03** | **0.05** | **0.05** | **0.01** |
| **M4S** | NA | NA | NA | NA | NA | NA | NA | NA | NA |
| **M5S** | **0.59** | **0.06** | **0.06** | **0.05** | **0.01** | **0.04** | **0.10** | **0.06** | **0.02** |

*^a^* see Supplementary Table S1.

*^b^* NA : not available; because most simulated test matrices did not contain any zero-valued cell, the *wNODF* method could not estimate their nestedness.

**Supplementary Table S3.** Type I errors of the *WINE* method for anti-nestedness estimation.

| **Anti-nestedness type I error, *WINE* method** | | | | | | | | | |
| --- | --- | --- | --- | --- | --- | --- | --- | --- | --- |
| **Test matrix** | **Null model** | | | | | | | | |
|  | **B** | **N** | **C1** | **R1** | **CR1** | **S** | **C2** | **R2** | **CR2** |
| **M1R** | **0.05***^a^* | **0** | **0.05** | **0.05** | **0** | **0.02** | **0.01** | **0.06** | **0** |
| **M2R** | **0.49** | **0.01** | **0.04** | **0.05** | **0** | **0** | **0.01** | **0.08** | **0** |
| **M3R** | **0.76** | **0** | **0** | 0.13 | **0** | **0** | **0** | **0.07** | **0** |
| **M4R** | **0.01** | **0** | **0** | **0** | **0** | **0** | **0** | **0.04** | **0** |
| **M5R** | **0** | **0** | **0** | **0.03** | **0** | **0** | **0** | **0.04** | **0** |
| **M1S** | 0.13 | **0.05** | **0.06** | **0.03** | **0** | **0.04** | **0.03** | **0.02** | **0** |
| **M2S** | **0.66** | **0.05** | **0.1** | **0.09** | **0** | **0.05** | **0.04** | **0.01** | **0** |
| **M3S** | **0.96** | **0.06** | **0.07** | **0.08** | **0** | **0.03** | **0.03** | **0.04** | **0** |
| **M4S** | 0.13 | **0.06** | **0.04** | **0.06** | **0.02** | **0.03** | **0.03** | **0.05** | **0** |
| **M5S** | **0.38** | **0.03** | **0.05** | **0.03** | **0.01** | **0.06** | **0.03** | **0.04** | **0** |

*^a^* For each test matrix pattern (table row) and each Null model pattern (table column), 100 test matrices were simulated, each of which being compared to 100 null-model matrices. This comparison provided a nestedness significance (p-value) for each simulated test matrix. The frequencies of nestedness p-values > 0.95 (F_0.95_) (*i.e.* of significant anti-nestedness) among the 100 simulated test matrices are indicated in the table. Favorable situations showing relatively few (F_0.95_ ≤ 0.10) false positive anti-nestedness estimations are in bold on grey cells and unfavorable situations showing relatively numerous (F_0.95_ ≥ 0.30) false positive anti-nestedness estimations are in white on black cells.

**Supplementary Table S4.** Type I errors of the *wNODF* method for anti-nestedness estimation.

| **Anti-nestedness type I error, *wNODF* method** | | | | | | | | | |
| --- | --- | --- | --- | --- | --- | --- | --- | --- | --- |
| **Test matrix** | **Null model** | | | | | | | | |
|  | **B** | **N** | **C1** | **R1** | **CR1** | **S** | **C2** | **R2** | **CR2** |
| **M1R** | **0.04***^a^* | **0.02** | **0.04** | **0.05** | **0** | **0** | **0** | **0.04** | **0** |
| **M2R** | **0.07** | **0.01** | **0.04** | **0.01** | **0** | **0.06** | **0.05** | **0.08** | **0.01** |
| **M3R** | **0** | **0.01** | **0** | **0.04** | **0** | **0.01** | **0.02** | **0.06** | **0** |
| **M4R** | NA*^b^* | NA | NA | NA | NA | NA | NA | NA | NA |
| **M5R** | **0** | **0.01** | **0** | **0.03** | **0** | **0** | **0** | **0.06** | **0** |
| **M1S** | **0.06** | **0.03** | **0.03** | **0.01** | **0** | **0.07** | **0.06** | **0.02** | **0** |
| **M2S** | **0.02** | **0.04** | **0.04** | **0.01** | **0** | **0.05** | **0.08** | **0.03** | **0** |
| **M3S** | **0** | **0.01** | **0.02** | **0.03** | **0** | **0.06** | **0.03** | **0.06** | **0** |
| **M4S** | NA | NA | NA | NA | NA | NA | NA | NA | NA |
| **M5S** | **0** | **0.05** | **0.05** | **0.04** | **0** | **0.04** | **0.03** | **0.02** | **0.01** |

*^a^* see Supplementary Table S3.

*^b^* NA: not available; because most simulated test matrices did not contain any zero-valued cell, the *wNODF* method could not estimate their nestedness.

**Supplementary Table S5.** Power of the *WINE* method for nestedness estimation.

| **Nestedness power, *WINE* method** | | | | | | | | | | |
| --- | --- | --- | --- | --- | --- | --- | --- | --- | --- | --- |
| **Test matrix** | **Nestedness score***^a^* | **Null model** | | | | | | | | |
|  |  | **B** | **N** | **C1** | **R1** | **CR1** | **S** | **C2** | **R2** | **CR2** |
| **M1** | 0.21 | 0.13*^b^* | **0.43** | **0.38** | **0.34** | 0.26 | **0.42** | **0.35** | **0.34** | **0.51** |
| **M2** | 0.27 | **0.01** | **0.54** | **0.40** | **0.35** | 0.18 | **0.56** | **0.43** | **0.49** | **0.67** |
| **M3** | 0.46 | **0** | **0.92** | **0.73** | **0.61** | **0.51** | **0.90** | **0.80** | **0.84** | **0.93** |
| **M4** | 0.57 | **0.63** | **0.77** | **0.7** | **0.66** | **0.59** | **0.79** | **0.58** | **0.70** | **0.76** |
| **M5** | 0.98 | **1** | **1** | **1** | **1** | **1** | **1** | **1** | **1** | **1** |

*^a^* Mean of nestedness scores of 100 simulated test matrices.

*^b^* For each test matrix pattern (table row) and each Null model pattern (table column), 100 test matrices were simulated, each of which being compared to 100 null-model matrices. This comparison provided a nestedness significance (p-value) for each simulated test matrix. The frequencies of nestedness p-values < 0.05 (F_0.05_) among the 100 simulated test matrices are indicated in the table. Favorable situations showing relatively few false negative nestedness estimations (F_0.05_ ≥ 0.30) are in bold on grey cells and unfavorable situations showing relatively numerous false negative nestedness estimations (F_0.05_ ≤ 0.10) are in white on black cells.

**Supplementary Table S6.** Power of the *wNODF* method for nestedness estimation.

| **Nestedness power, *wNODF* method** | | | | | | | | | | |
| --- | --- | --- | --- | --- | --- | --- | --- | --- | --- | --- |
| **Test matrix** | **Mean estimate***^a^* | **Null model** | | | | | | | | |
|  |  | **B** | **N** | **C1** | **R1** | **CR1** | **S** | **C2** | **R2** | **CR2** |
| **M1** | 26.4 | **0.03***^b^* | **0.40** | **0.35** | 0.22 | **0.09** | **0.34** | 0.27 | 0.23 | **0.38** |
| **M2** | 34.1 | **0.01** | **0.53** | **0.36** | **0.35** | 0.13 | **0.44** | **0.36** | **0.38** | **0.61** |
| **M3** | 38.5 | 0.14 | **0.77** | **0.54** | **0.50** | 0.25 | **0.73** | **0.60** | **0.59** | **0.80** |
| **M4** | NA*^c^* | NA | NA | NA | NA | NA | NA | NA | NA | NA |
| **M5** | 42.9 | **0.6** | **1** | **0.99** | **0.99** | **0.98** | **0.99** | **0.99** | **0.98** | **0.99** |

*^a^*^,^*^b^* see Supplementary Table S5.

*^c^*NA: not available; because most simulated test matrices did not contain any zero-valued cell, the *wNODF* method could not estimate their nestedness.

**Supplementary Table S7.** Type I errors of the *edge betweenness* method for modularity estimation.

| **Modularity type I error, *edge betweenness* method** | | | | | | | | | |
| --- | --- | --- | --- | --- | --- | --- | --- | --- | --- |
| **Test matrix** | **Null model** | | | | | | | | |
|  | **B** | **N** | **C1** | **R1** | **CR1** | **S** | **C2** | **R2** | **CR2** |
| **M1R** | 0.16*^a^* | **0.01** | **0.01** | **0.05** | **0** | **0** | **0** | **0.02** | **0** |
| **M2R** | **0.46** | **0.10** | **0.10** | 0.24 | **0.10** | **0.03** | **0.02** | **0.02** | **0.02** |
| **M3R** | **0.33** | **0.10** | **0.08** | 0.25 | **0.08** | **0** | **0** | **0.02** | **0** |
| **M4R** | **0** | **0.01** | **0** | **0** | **0** | **0.01** | **0.02** | **0.07** | **0.02** |
| **M5R** | **0** | **0** | **0** | **0.03** | **0** | **0** | **0** | **0.05** | **0** |
| **M1S** | 0.16 | **0.05** | **0.06** | **0.05** | **0.03** | **0.06** | **0.05** | **0.04** | **0.04** |
| **M2S** | **0.42** | 0.19 | 0.17 | 0.17 | 0.13 | **0.04** | **0.04** | **0.05** | **0.03** |
| **M3S** | **0.79** | **0.48** | **0.59** | **0.55** | **0.49** | **0.02** | **0** | **0.03** | **0** |
| **M4S** | **0.02** | **0.06** | **0.05** | **0.06** | **0.05** | **0.04** | **0.03** | **0.04** | **0.02** |
| **M5S** | **0.06** | **0.08** | **0.05** | **0.07** | **0.04** | **0.03** | **0.03** | **0.02** | **0.02** |

*^a^* For each test matrix pattern (table row) and each null model pattern (table column), 100 test matrices were simulated, each of which being compared to 100 null-model matrices. This comparison provided a modularity significance (p-value) for each simulated test matrix. The frequencies of modularity p-values < 0.05 (F_0.05_) among the 100 simulated test matrices are indicated in the table. Favorable situations showing relatively few (F_0.05_ ≤ 0.10) false positive modularity estimations are in bold on grey cells and unfavorable situations showing relatively numerous (F_0.05_ ≥ 0.30) false positive modularity estimations are in white on black cells.

**Supplementary Table S8.** Type I errors of the *fast greedy* method for modularity estimation.

| **Modularity type I error, *fast greedy* method** | | | | | | | | | |
| --- | --- | --- | --- | --- | --- | --- | --- | --- | --- |
| **Test matrix** | **Null model** | | | | | | | | |
|  | **B** | **N** | **C1** | **R1** | **CR1** | **S** | **C2** | **R2** | **CR2** |
| **M1R** | **0.01***^a^* | **0.03** | **0.02** | **0.02** | **0** | **0.01** | **0.02** | **0.03** | **0.01** |
| **M2R** | **0.72** | **0.33** | **0.31** | **0.49** | **0.30** | **0.01** | **0** | **0.05** | **0** |
| **M3R** | **0.60** | 0.27 | 0.26 | **0.44** | 0.15 | **0** | **0** | **0.09** | **0** |
| **M4R** | **0** | **0** | **0** | **0** | **0** | **0.01** | **0.01** | **0.03** | **0.01** |
| **M5R** | **0** | **0** | **0** | **0** | **0** | **0** | **0** | **0.04** | **0** |
| **M1S** | **0.10** | **0.06** | **0.03** | **0.04** | **0.02** | **0.03** | **0.04** | **0.05** | **0.02** |
| **M2S** | **0.88** | **0.64** | **0.65** | **0.64** | **0.61** | **0.05** | **0.04** | **0.05** | **0.02** |
| **M3S** | **0.90** | **0.64** | **0.69** | **0.68** | **0.64** | **0.03** | **0.02** | **0.04** | **0.02** |
| **M4S** | **0** | **0** | **0** | **0** | **0** | **0.05** | **0.02** | **0.03** | **0.02** |
| **M5S** | **0.02** | **0.07** | **0.09** | **0.08** | **0.06** | **0.02** | **0.03** | **0.01** | **0.01** |

*^a^* see Supplementary Table S7.

**Supplementary Table S9.** Type I errors of the *louvain* method for modularity estimation.

| **Modularity type I error, *louvain* method** | | | | | | | | | |
| --- | --- | --- | --- | --- | --- | --- | --- | --- | --- |
| **Test matrix** | **Null model** | | | | | | | | |
|  | **B** | **N** | **C1** | **R1** | **CR1** | **S** | **C2** | **R2** | **CR2** |
| **M1R** | **0.02***^a^* | **0** | **0** | **0.01** | **0** | **0** | **0** | **0** | **0** |
| **M2R** | **0.45** | 0.22 | 0.18 | **0.34** | 0.18 | **0.02** | **0** | **0.03** | **0** |
| **M3R** | 0.14 | **0.02** | **0.01** | **0.05** | **0.01** | **0** | **0** | **0** | **0** |
| **M4R** | **0** | **0** | **0** | **0** | **0** | **0** | **0** | **0** | **0** |
| **M5R** | **0** | **0** | **0** | **0** | **0** | **0** | **0** | **0** | **0** |
| **M1S** | **0.03** | **0.01** | **0.01** | **0.02** | **0.01** | **0.01** | **0.02** | **0.01** | **0.01** |
| **M2S** | **0.66** | **0.31** | 0.29 | **0.30** | 0.23 | **0.02** | **0.01** | **0.01** | **0** |
| **M3S** | **0.64** | **0.34** | **0.30** | **0.35** | 0.27 | **0** | **0** | **0.01** | **0** |
| **M4S** | **0** | **0** | **0** | **0** | **0** | **0** | **0** | **0** | **0** |
| **M5S** | **0.01** | **0.01** | **0.01** | **0.01** | **0.01** | **0.01** | **0** | **0** | **0** |

*^a^* see Supplementary Table S7.

**Supplementary Table S10.** Type I errors of the *spinglass* method for modularity estimation.

| **Modularity type I error, *spinglass* method** | | | | | | | | | |
| --- | --- | --- | --- | --- | --- | --- | --- | --- | --- |
| **Test matrix** | **Null model** | | | | | | | | |
|  | **B** | **N** | **C1** | **R1** | **CR1** | **S** | **C2** | **R2** | **CR2** |
| **M1R** | **0***^a^* | **0** | **0** | **0** | **0** | **0.01** | **0** | **0** | **0** |
| **M2R** | **0** | **0** | **0** | **0** | **0.01** | **0** | **0.01** | **0** | **0** |
| **M3R** | **0** | **0** | **0** | **0** | **0** | **0** | **0** | **0** | **0** |
| **M4R** | **0** | **0** | **0** | **0** | **0** | **0** | **0** | **0** | **0** |
| **M5R** | **0** | **0** | **0** | **0** | **0** | **0** | **0** | **0** | **0** |
| **M1S** | **0** | **0** | **0** | **0** | **0** | NA*^b^* | NA | NA | NA |
| **M2S** | **0.01** | **0** | **0** | **0** | **0** | **0** | **0** | **0** | **0** |
| **M3S** | **0.01** | **0** | **0** | **0** | **0** | **0** | **0** | **0** | **0** |
| **M4S** | **0** | **0** | **0** | **0** | **0** | **0** | **0** | **0** | **0** |
| **M5S** | **0.01** | **0** | **0** | **0** | **0** | **0** | **0** | **0** | **0** |

*^a^* see Supplementary Table S7.

*^b^* NA: not available; many null-model matrices had rows and/or columns entirely made of zero-valued cells and the *spinglass* method could not estimate their modularity.

**Supplementary Table S11.** Type I errors of the *edge betweenness* method for anti-modularity estimation.

| **Anti-modularity type I error, *edge betweenness* method** | | | | | | | | | |
| --- | --- | --- | --- | --- | --- | --- | --- | --- | --- |
| **Test matrix** | **Null model** | | | | | | | | |
|  | **B** | **N** | **C1** | **R1** | **CR1** | **S** | **C2** | **R2** | **CR2** |
| **M1R** | **0***^a^* | **0.02** | **0.02** | **0.02** | **0.01** | **0.04** | **0.07** | **0.01** | **0.01** |
| **M2R** | **0** | **0** | **0.02** | **0** | **0** | **0.03** | **0.03** | **0.01** | **0** |
| **M3R** | **0** | **0** | **0.01** | **0** | **0** | 0.21 | **0.34** | **0.03** | **0.03** |
| **M4R** | **0** | **0.02** | **0.02** | **0** | **0** | 0.12 | 0.14 | **0.04** | **0.04** |
| **M5R** | **0.02** | 0.13 | 0.12 | **0.01** | **0.01** | **0.33** | **0.49** | **0** | **0** |
| **M1S** | **0** | **0.03** | **0.04** | **0.03** | **0.02** | **0.02** | **0.02** | **0.02** | **0.01** |
| **M2S** | **0** | **0.05** | **0.04** | **0.03** | **0.03** | **0.03** | **0.03** | **0.02** | **0** |
| **M3S** | **0** | **0** | **0** | **0** | **0** | **0.01** | **0.01** | **0.02** | **0.01** |
| **M4S** | **0.02** | **0** | **0** | **0** | **0** | **0.01** | **0.01** | **0.03** | **0** |
| **M5S** | **0** | **0** | **0** | **0** | **0** | **0.04** | **0.02** | **0.03** | **0.02** |

*^a^* For each test matrix pattern (table row) and each Null model pattern (table column), 100 test matrices were simulated, each of which being compared to 100 null-model matrices. This comparison provided a modularity significance (p-value) for each simulated test matrix. The frequencies of modularity p-values > 95 (F_0.95_) (*i.e.* of significant anti-modularity) among the 100 simulated test matrices are indicated in the table. Favorable situations showing relatively few (F_0.95_ ≤ 0.10) false positive anti-modularity estimations are in bold on grey cells and unfavorable situations showing relatively numerous (F_0.95_ ≥ 0.30) false positive anti-modularity estimations are in white on black cells.

**Supplementary Table S12.** Type I errors of the *fast greedy* method for anti-modularity estimation.

| **Anti-modularity type I error, *fast greedy* method** | | | | | | | | | |
| --- | --- | --- | --- | --- | --- | --- | --- | --- | --- |
| **Test matrix** | **Null model** | | | | | | | | |
|  | **B** | **N** | **C1** | **R1** | **CR1** | **S** | **C2** | **R2** | **CR2** |
| **M1R** | **0.05***^a^* | 0.17 | 0.19 | **0.09** | **0.04** | 0.20 | 0.22 | **0.05** | **0.04** |
| **M2R** | **0** | **0.01** | **0.01** | **0.01** | **0** | 0.29 | **0.32** | **0.07** | **0.06** |
| **M3R** | **0** | **0** | **0.01** | **0** | **0** | **0.42** | **0.54** | **0.02** | **0.02** |
| **M4R** | **0.82** | **0.65** | **0.64** | **0.64** | **0.53** | **0.08** | **0.08** | **0.04** | **0.03** |
| **M5R** | **0.58** | **0.61** | **0.65** | **0.34** | **0.33** | **0.88** | **0.96** | **0.06** | **0.06** |
| **M1S** | **0** | **0.04** | **0.05** | **0.04** | **0.04** | **0.09** | **0.04** | **0.05** | **0.02** |
| **M2S** | **0** | **0** | **0** | **0** | **0** | 0.11 | **0.09** | **0.08** | **0.03** |
| **M3S** | **0** | **0.01** | **0.01** | **0** | **0** | **0.06** | **0.07** | **0.07** | **0.02** |
| **M4S** | **0.71** | **0.46** | **0.45** | **0.47** | **0.40** | **0.05** | **0.05** | **0.04** | **0.04** |
| **M5S** | **0.01** | **0.01** | **0.01** | **0.01** | **0.01** | **0.08** | **0.05** | **0.08** | **0.04** |

*^a^* see Supplementary Table S11.

**Supplementary Table S13.** Type I errors of the *louvain* method for anti-modularity estimation.

| **Anti-modularity type I error, *louvain* method** | | | | | | | | | |
| --- | --- | --- | --- | --- | --- | --- | --- | --- | --- |
| **Test matrix** | **Null model** | | | | | | | | |
|  | **B** | **N** | **C1** | **R1** | **CR1** | **S** | **C2** | **R2** | **CR2** |
| **M1R** | **0.01***^a^* | **0.08** | **0.06** | **0.01** | **0.01** | **0.05** | **0.10** | **0.01** | **0** |
| **M2R** | **0** | **0** | **0** | **0** | **0** | 0.14 | 0.17 | **0.01** | **0.01** |
| **M3R** | **0** | **0.01** | **0** | **0** | **0** | 0.12 | 0.20 | **0** | **0** |
| **M4R** | **0.52** | 0.26 | **0.31** | 0.25 | 0.22 | **0** | **0** | **0** | **0** |
| **M5R** | 0.26 | 0.27 | 0.25 | **0.06** | **0.06** | **0.74** | **0.85** | **0** | **0** |
| **M1S** | **0** | **0** | **0** | **0** | **0** | **0.01** | **0.02** | **0** | **0** |
| **M2S** | **0** | **0** | **0** | **0** | **0** | **0.02** | **0.02** | **0.01** | **0** |
| **M3S** | **0** | **0** | **0** | **0** | **0** | **0** | **0** | **0** | **0** |
| **M4S** | **0.40** | 0.16 | 0.18 | 0.19 | 0.13 | **0** | **0** | **0** | **0** |
| **M5S** | **0** | **0** | **0** | **0** | **0** | **0** | **0** | **0** | **0** |

*^a^* see Supplementary Table S11.

**Supplementary Table S14.** Type I errors of the *spinglass* method for anti-modularity estimation.

| **Anti-modularity type I error, *spinglass* method** | | | | | | | | | |
| --- | --- | --- | --- | --- | --- | --- | --- | --- | --- |
| **Test matrix** | **Null model** | | | | | | | | |
|  | **B** | **N** | **C1** | **R1** | **CR1** | **S** | **C2** | **R2** | **CR2** |
| **M1R** | **0***^a^* | **0** | **0** | **0** | **0** | **0** | **0** | **0** | **0** |
| **M2R** | **0** | **0.01** | **0.01** | **0.01** | **0** | **0** | **0** | **0** | **0** |
| **M3R** | **0** | **0** | **0** | **0** | **0** | **0** | **0** | **0** | **0** |
| **M4R** | **0** | **0** | **0** | **0** | **0** | **0** | **0** | **0** | **0** |
| **M5R** | **0** | **0** | **0** | **0** | **0** | **0** | **0** | **0** | **0** |
| **M1S** | **0** | **0** | **0** | **0** | **0** | NA*^b^* | NA | NA | NA |
| **M2S** | **0** | **0** | **0** | **0** | **0** | **0** | **0** | **0** | **0** |
| **M3S** | **0** | **0** | **0** | **0** | **0** | **0.01** | **0** | **0** | **0** |
| **M4S** | **0** | **0** | **0** | **0** | **0** | **0** | **0** | **0** | **0** |
| **M5S** | **0** | **0** | **0** | **0** | **0** | **0** | **0** | **0** | **0** |

*^a^* see Supplementary Table S11.

*^b^* NA: not available; many null-model matrices had rows and/or columns entirely made of zero-valued cells and the *spinglass* method could not estimate their modularity.

**Supplementary Table S15.** Power of the *edge betweenness* method for modularity estimation.

| **Modularity power, *edge betweenness* method** | | | | | | | | | | |
| --- | --- | --- | --- | --- | --- | --- | --- | --- | --- | --- |
| **Test matrix** | **Modularity score***^a^* | **Null model** | | | | | | | | |
|  |  | **B** | **N** | **C1** | **R1** | **CR1** | **S** | **C2** | **R2** | **CR2** |
| **M6** | 0.080 | **0.47***^b^* | **0.36** | **0.36** | **0.39** | **0.36** | 0.22 | 0.22 | 0.23 | 0.19 |
| **M7** | 0.084 | **0.48** | **0.35** | **0.36** | **0.38** | **0.33** | 0.14 | 0.12 | 0.11 | **0.09** |
| **M8** | 0.101 | **0.32** | 0.11 | 0.15 | 0.11 | 0.10 | **0.08** | **0.10** | **0.08** | **0.08** |
| **M9** | 0.112 | **0.49** | **0.33** | **0.31** | **0.31** | 0.28 | **0.03** | **0.03** | **0.02** | **0.02** |
| **M10** | 0.195 | **0.93** | **0.91** | **0.91** | **0.90** | **0.90** | **0.61** | **0.58** | **0.56** | **0.56** |
| **M11** | 0.234 | **0.98** | **0.98** | **0.98** | **0.98** | **0.98** | **0.86** | **0.86** | **0.85** | **0.85** |

*^a^* Mean of modularity scores of 100 simulated test matrices.

*^b^* For each test matrix pattern (table row) and each Null model pattern (table column), 100 test matrices were simulated, each of which being compared to 100 null-model matrices. This comparison provided a modularity significance (p-value) for each simulated test matrix. The frequencies of modularity p-values < 0.05 (F_0.05_) among the 100 simulated test matrices are indicated in the table. Favorable situations showing relatively few false negative modularity estimations (F_0.05_ ≥ 0.30) are in bold on grey cells and unfavorable situations showing relatively numerous false negative modularity estimations (F_0.05_ ≤ 0.10) are in white on black cells.

**Supplementary Table S16.** Power of the *fast greedy* method for modularity estimation.

| **Modularity power, *fast greedy* method** | | | | | | | | | | |
| --- | --- | --- | --- | --- | --- | --- | --- | --- | --- | --- |
| **Test matrix** | **Modularity score***^a^* | **Null model** | | | | | | | | |
|  |  | **B** | **N** | **C1** | **R1** | **CR1** | **S** | **C2** | **R2** | **CR2** |
| **M6** | 0.097 | 0.15*^b^* | **0.09** | **0.10** | 0.11 | **0.09** | **0.37** | **0.36** | **0.42** | **0.32** |
| **M7** | 0.108 | **0.33** | 0.15 | 0.18 | 0.16 | 0.13 | 0.13 | 0.13 | 0.13 | 0.11 |
| **M8** | 0.219 | **1** | **0.99** | **1** | **0.98** | **0.98** | **0.55** | **0.50** | **0.50** | **0.40** |
| **M9** | 0.212 | **1** | **0.99** | **1** | **1** | **1** | **0.63** | **0.57** | **0.61** | **0.52** |
| **M10** | 0.294 | **1** | **1** | **1** | **1** | **1** | **1** | **1** | **1** | **1** |
| **M11** | 0.278 | **1** | **1** | **1** | **1** | **1** | **1** | **1** | **1** | **1** |

*^a^*^,^*^b^* see Supplementary Table S15.

**Supplementary Table S17.** Power of the *louvain* method for modularity estimation.

| **Modularity power, *louvain* method** | | | | | | | | | | |
| --- | --- | --- | --- | --- | --- | --- | --- | --- | --- | --- |
| **Test matrix** | **Modularity score***^a^* | **Null model** | | | | | | | | |
|  |  | **B** | **N** | **C1** | **R1** | **CR1** | **S** | **C2** | **R2** | **CR2** |
| **M6** | 0.097 | **0.02***^b^* | **0** | **0** | **0** | **0** | **0.02** | **0.01** | **0.02** | **0.01** |
| **M7** | 0.106 | **0.04** | **0.01** | **0.01** | **0** | **0** | **0.01** | **0.01** | **0.01** | **0.01** |
| **M8** | 0.213 | **0.98** | **0.94** | **0.96** | **0.95** | **0.95** | **0.35** | **0.34** | 0.29 | 0.24 |
| **M9** | 0.211 | **1** | **0.99** | **1** | **1** | **1** | **0.41** | **0.34** | **0.36** | 0.29 |
| **M10** | 0.295 | **1** | **1** | **1** | **1** | **1** | **1** | **1** | **1** | **1** |
| **M11** | 0.278 | **1** | **1** | **1** | **1** | **1** | **1** | **1** | **1** | **1** |

*^a^*^,^*^b^* see Supplementary Table S15.

**Supplementary Table S18.** Power of the *spinglass* method for modularity estimation.

| **Modularity power, *spinglass* method** | | | | | | | | | | |
| --- | --- | --- | --- | --- | --- | --- | --- | --- | --- | --- |
| **Test matrix** | **Modularity score***^a^* | **Null model** | | | | | | | | |
|  |  | **B** | **N** | **C1** | **R1** | **CR1** | **S** | **C2** | **R2** | **CR2** |
| **M6** | 0.060 | **0.38***^b^* | 0.28 | 0.28 | 0.28 | 0.25 | 0.20 | 0.20 | 0.20 | 0.17 |
| **M7** | 0.049 | **0.03** | **0.01** | **0.01** | **0.01** | **0.01** | **0** | **0** | **0** | **0** |
| **M8** | 0.065 | **0** | **0** | **0** | **0** | **0** | **0** | **0** | **0.01** | **0** |
| **M9** | 0.067 | **0.02** | **0.01** | **0.01** | **0.01** | **0.01** | **0.02** | **0.02** | **0.01** | **0.01** |
| **M10** | 0.072 | **0.11** | **0.01** | **0** | **0** | **0** | **0.06** | **0.05** | **0.05** | **0.03** |
| **M11** | 0.091 | **1** | **1** | **1** | **1** | **1** | **1** | **1** | **1** | **1** |

*^a^*^,^*^b^* see Supplementary Table S15.

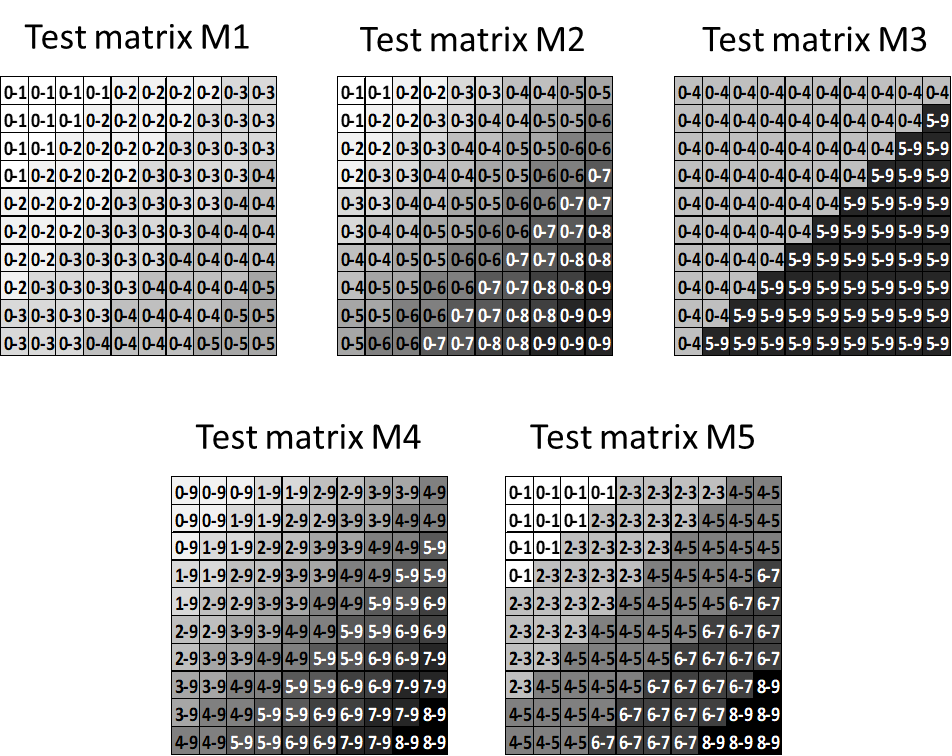

**Supplementary Figure S1.** Patterns of test matrices used to evaluate the statistical power of nestedness evaluation methods. A test matrix is simulated by sampling randomly, for each cell, an integer value in the indicated interval.

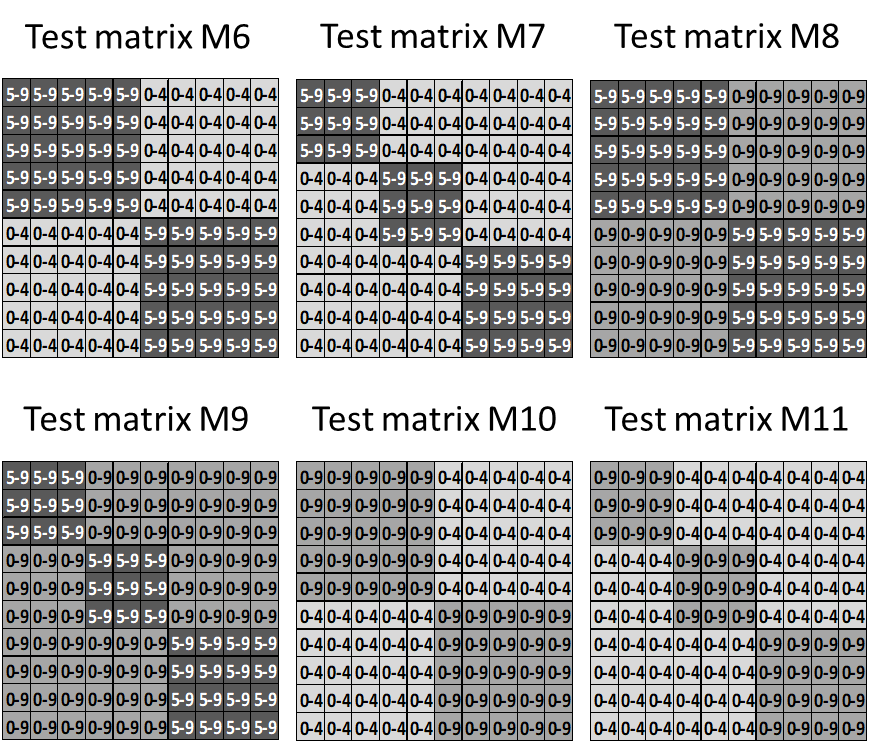

**Supplementary Figure S2.** Patterns of test matrices used to evaluate the statistical power of modularity evaluation methods. A test matrix is simulated by sampling randomly, for each cell, an integer value in the indicated interval.
