## Supplementary material for "The quasi-universality of nestedness in the structure of quantitative plant-parasite interactions": Suppl Methods 1 and 2; Suppl. Tables 19 to 22: Table S19.docx

**Supplementary Table S19.** Analysis of nestnedness of plant-parasite interaction matrices with the *wNODF* method.

| **Matrix number** | **Nestedness score** | **Null model***^a^* | | | | | | |
| --- | --- | --- | --- | --- | --- | --- | --- | --- |
|  |  | **B** | **N** | **C1** | **R1** | **S** | **C2** | **R2** |
| **1** | 30.9 | **0.05***^b^* | **0** | **0** | **0** | **0** | **0** | **0** |
| **2** | 43.7 | 0.06 | **0** | **0** | **0** | **0** | **0** | **0** |
| **3** | 47.6 | **0** | **0** | **0** | **0** | **0** | **0** | **0** |
| **4** | 46.9 | **0** | **0** | **0** | **0** | **0** | **0** | **0** |
| **5** | 23.2 | 0.24 | **0** | 0.42 | **0** | **0** | 0.36 | **0** |
| **6** | 55.4 | 0.15 | **0** | **0** | **0** | **0** | **0** | **0** |
| **7** | 43.9 | 0.41 | **0** | **0** | **0** | **0** | **0** | **0** |
| **8** | 49.4 | 0.66 | **0** | **0** | **0** | **0** | **0** | **0** |
| **9** | 42.4 | **0.01** | **0** | **0** | **0** | **0** | **0** | **0** |
| **10** | 72.7 | **0** | **0** | **0** | **0** | **0** | **0** | **0** |
| **11** | 75.4 | **0** | **0** | **0** | **0** | **0** | **0** | **0** |
| **12** | 10.1 | 0.68 | 0.85 | 0.67 | **1** | 0.73 | 0.54 | 0.90 |
| **13** | 30.0 | 0.16 | 0.38 | 0.48 | 0.07 | 0.35 | 0.39 | 0.52 |
| **14** | 22.3 | 0.76 | **1** | **1** | **1** | 0.06 | **1** | 0.22 |
| **15** | 41.9 | 0.31 | **0** | **0** | **0** | **0** | **1** | **0** |
| **16** | 38.6 | 0.09 | **0** | **0** | **0** | **0** | **0** | **0** |
| **17** | 51.0 | **0** | **0** | **0** | **0** | **0** | **0** | **0** |
| **18** | 59.2 | **0** | **0** | **0** | **0** | **0** | **0** | **0** |
| **19** | 9.4 | 0.89 | **0.01** | **0.01** | 0.80 | **0.04** | **0** | **0.01** |
| **20** | 15.0 | 0.98 | **0** | **0** | **0** | **0** | 0.12 | **0** |
| **21** | 27.6 | 0.33 | 0.42 | 0.68 | **0** | 0.51 | **1** | 0.59 |
| **22** | 49.2 | **0** | **0** | **0** | **0** | **0** | **0** | **0** |
| **23** | 6.1 | 0.65 | 0.82 | 0.28 | **1** | 0.82 | 0.48 | 0.85 |
| **24** | 22.4 | 0.29 | **0.98** | **0.96** | 0.72 | **0.95** | 0.93 | 0.72 |
| **25** | 44.1 | 0.20 | **0** | **0** | **0** | **0** | **0.02** | **0** |
| **26** | 34.3 | **0** | **0** | **0** | **0** | **0** | **0** | **0** |
| **27** | 51.2 | 0.41 | **0** | **0** | **0.01** | **0** | **0.01** | **0** |
| **28** | 58.0 | 0.27 | **0** | **0** | **0** | **0** | **0** | **0** |
| **29** | 22.3 | **0.97** | 0.81 | 0.62 | **0.02** | 0.88 | **0.96** | 0.18 |
| **30** | 66.8 | **0.01** | **0** | **0** | **0** | **0** | **0** | **0** |
| **31** | 21.3 | **0.05** | 0.30 | 0.49 | **0.04** | 0.21 | 0.35 | 0.06 |
| **32** | 11.1 | 0.13 | 0.65 | 0.55 | 0.90 | 0.62 | 0.51 | 0.74 |

*^a^* See Supplementary Methods 2 for details of the null models.

*^b^* Nestedness significance: the probability value (p-value) indicates the frequency of null-model matrices showing a strictly higher nestedness score than that of the actual matrix. P-values ≤ 0.05 (significant nestedness) are in bold on grey cells and p-values > 0.95 (significant anti-nestedness) are in white on black cells.
