## Supplementary material for "The quasi-universality of nestedness in the structure of quantitative plant-parasite interactions": Suppl Methods 1 and 2; Suppl. Tables 19 to 22: Table S21.docx

**Supplementary Table S21.** Pearson’s coefficients of correlation between parasites host range breadth and pathogenicity or between plant resistance efficiency and scope in the 32 analysed matrices for different thresholds separating hosts and non-hosts (or parasites included or not included in the resistance scope). Each threshold (from 10% to 90%) corresponds to a percentage of the maximal pathogenicity value in each matrix. For some thresholds and some matrices, the coefficient of correlation could not be calculated because too few data remained (NA).

| **Matrix** | **Threshold** | | | | | | | | |
| --- | --- | --- | --- | --- | --- | --- | --- | --- | --- |
|  | **10%** | **20%** | **30%** | **40%** | **50%** | **60%** | **70%** | **80%** | **90%** |
| **Parasites host range breadth and pathogenicity** | | | | | | | | | |
| **1** | 0.85***^a^* | 0.83** | 0.77* | 0.76ns | 0.65ns | -0.20ns | 0.86ns | NA | NA |
| **2** | 0.91** | 0.89* | 0.79ns | 0.83ns | 1.00** | 0.94ns | 1.00ns | NA | NA |
| **3** | 0.68* | 0.04ns | 0.03ns | 0.17ns | 0.31ns | 0.12ns | 0.05ns | 0.08ns | NA |
| **4** | 0.77* | 0.23ns | 0.09ns | -0.41ns | -0.37ns | -0.33ns | NA | NA | NA |
| **5** | 0.18ns | 0.11ns | 0.05ns | -0.30ns | -0.59* | -0.60* | -0.27ns | 0.22ns | 0.91*** |
| **6** | 0.79*** | 0.56* | 0.86*** | 0.85*** | 0.61ns | 0.77** | 0.35ns | 0.56ns | NA |
| **7** | 0.72** | 0.02ns | 0.17ns | 0.05ns | -0.03ns | -0.36ns | NA | NA | NA |
| **8** | 0.38ns | 0.44* | 0.49* | 0.49* | 0.47* | 0.45ns | 0.38ns | 0.48ns | 0.09ns |
| **9** | 0.40*** | 0.55*** | 0.69*** | 0.51*** | 0.55*** | 0.31* | 0.62* | -0.10ns | 0.11ns |
| **10** | 0.86*** | 0.55** | 0.57** | 0.18ns | -0.23ns | -0.43* | -0.34ns | -0.40* | -0.23ns |
| **11** | 0.59*** | 0.75*** | 0.22ns | -0.22ns | -0.37ns | -0.44* | -0.53** | -0.36ns | -0.24ns |
| **12** | 0.57* | 0.55* | 0.52* | 0.46ns | 0.67** | 0.70** | 0.75** | 0.21ns | 0.15ns |
| **13** | 0.43 | 0.57* | 0.48ns | 0.61* | 0.84*** | 0.85*** | 0.78** | -0.08ns | -0.17ns |
| **14** | 0.19 | 0.13ns | 0.20* | 0.20ns | -0.04ns | -0.06ns | 0.19ns | 0.28** | 0.32** |
| **15** | 0.15 | 0.31** | 0.37*** | 0.35*** | 0.40*** | 0.31** | 0.38*** | 0.42** | 0.67** |
| **16** | NA | 0.68ns | 0.55ns | 0.64ns | 0.51ns | 0.69ns | 0.75ns | 0.85ns | NA |
| **17** | 0.44 | 0.63ns | 0.69ns | 0.76ns | 0.91* | 0.96* | 0.88ns | 0.18ns | NA |
| **18** | 0.43 | -0.68ns | -0.78* | -0.66ns | -0.72ns | -0.73ns | NA | NA | NA |
| **19** | -0.67*** | -0.28ns | 0.38* | -0.44* | -0.43* | -0.22ns | NA | NA | NA |
| **20** | 0.51*** | 0.62*** | 0.29ns | 0.05ns | 0.20ns | 0.33ns | 0.00ns | 0.17ns | NA |
| **21** | -0.50** | -0.56*** | -0.36* | -0.12ns | 0.21ns | -0.09ns | -0.03ns | -0.14ns | NA |
| **22** | 0.27ns | 0.31ns | 0.07ns | -0.08ns | -0.28ns | 0.09ns | 0.23ns | 0.38ns | -0.19ns |
| **23** | 0.53ns | -0.20ns | -0.06ns | -0.01ns | -0.03ns | -0.25ns | -0.20ns | 1.00*** | NA |
| **24** | -0.25ns | -0.22ns | -0.15ns | -0.08ns | -0.29ns | -0.16ns | 0.33ns | 0.96** | NA |
| **25** | 0.14ns | 0.25ns | 0.06ns | 0.35ns | 0.46ns | 0.90ns | NA | NA | NA |
| **26** | 0.24ns | 0.58* | 0.67* | 0.61* | 0.57* | 0.73** | 0.75** | 0.68* | 0.77* |
| **27** | -0.02ns | 0.64ns | 0.71* | 0.50ns | 0.66ns | 0.95** | 0.58ns | 1.00* | NA |
| **28** | 0.44ns | 0.89** | 0.70* | 0.82* | 0.66ns | -0.34ns | -0.51ns | -0.18ns | NA |
| **29** | 0.24ns | -0.34ns | 0.01ns | -0.31ns | 0.18ns | 0.42ns | 0.98ns | NA | NA |
| **30** | 0.86* | 0.69ns | 0.77ns | 0.94* | 0.94ns | NA | NA | NA | NA |
| **31** | 0.37ns | 0.37ns | 0.58ns | 0.56ns | 0.53ns | -0.39ns | 0.05ns | 0.92** | NA |
| **32** | 0.28ns | 0.28ns | -0.04ns | 0.10ns | 0.44ns | 0.79* | 0.86* | NA | NA |
| **Plant resistance efficiency and scope** | | | | | | | | | |
| **1** | 0.70*** | 0.58** | 0.38ns | 0.48* | 0.61** | 0.39ns | 0.24ns | 0.27ns | 0.30ns |
| **2** | -0.09ns | -0.01ns | -0.14ns | 0.11ns | 0.01ns | -0.04ns | -0.13ns | 0.35ns | 0.11ns |
| **3** | 0.60** | 0.50* | 0.50* | 0.39ns | 0.48* | 0.68*** | 0.53* | 0.41ns | 0.18ns |
| **4** | 0.07ns | 0.19ns | 0.23ns | 0.36ns | 0.36ns | 0.39ns | 0.43ns | 0.61** | 0.61** |
| **5** | NA | -0.69ns | -0.63ns | 0.70ns | 0.76ns | 0.81* | 0.87** | 0.92*** | 0.79** |
| **6** | -0.56ns | -0.67* | -0.55ns | -0.35ns | -0.28ns | -0.55ns | 0.46ns | 0.52ns | 0.50ns |
| **7** | 0.32ns | 0.27ns | 0.78** | 0.76** | 0.74** | 0.72** | 0.11ns | 0.52ns | 0.02ns |
| **8** | 0.58ns | 0.23ns | 0.03ns | 0.67ns | 0.52ns | 0.46ns | 0.85** | 0.79* | 0.56ns |
| **9** | 0.35ns | -0.22ns | 0.71** | 0.73** | 0.83*** | 0.85*** | -0.06ns | 0.10ns | -0.11ns |
| **10** | 0.92*** | 0.79*** | 0.85*** | 0.92*** | 0.94*** | 0.89*** | 0.81*** | 0.85*** | 0.73*** |
| **11** | 0.43ns | 0.48ns | 0.88*** | 0.92*** | 0.92*** | 0.90*** | 0.92*** | 0.79*** | 0.70*** |
| **12** | NA | NA | -0.53ns | -0.71* | -0.63* | 0.59* | 0.74** | 0.26ns | 0.11ns |
| **13** | 0.01ns | -0.34ns | -0.58ns | -0.64* | -0.36ns | 0.41ns | 0.61* | 0.54ns | 0.31ns |
| **14** | 0.65ns | 0.40ns | 0.34ns | 0.40ns | 0.28ns | 0.31ns | 0.38ns | 0.31ns | 0.61* |
| **15** | 0.24ns | 0.61* | 0.67** | 0.72** | 0.75** | 0.75** | 0.76*** | 0.82*** | 0.73** |
| **16** | NA | NA | 0.44ns | 0.52ns | 0.30ns | 0.94ns | 0.97*** | 0.82** | 0.46ns |
| **17** | 0.03ns | 0.07ns | 0.41** | 0.61*** | 0.56*** | 0.75*** | 0.63*** | 0.64*** | 0.44*** |
| **18** | 0.75ns | 0.88** | 0.40ns | 0.33ns | 0.41ns | 0.31ns | NA | NA | 0.95*** |
| **19** | 0.29ns | 0.18ns | 0.08ns | 0.60ns | 0.70ns | 0.61ns | 0.63ns | 0.70ns | 0.70ns |
| **20** | 0.73* | 0.71* | 0.73* | 0.73* | 0.71* | 0.65* | 0.66* | 0.57ns | 0.59ns |
| **21** | -0.58ns | 0.07ns | 0.51ns | 0.75ns | 0.77* | 0.78* | 0.85** | 0.77* | 0.75* |
| **22** | 0.97** | 0.68ns | 0.90** | 0.94*** | 0.84** | 0.83** | 0.85** | 0.85** | 0.84** |
| **23** | NA | NA | -0.18ns | -0.14ns | -0.57ns | -0.70* | 0.60* | 0.65* | 0.44ns |
| **24** | NA | -1.00*** | -0.94*** | -0.73* | -0.53ns | -0.44ns | -0.43ns | 0.83*** | -0.19ns |
| **25** | 0.58ns | 0.82** | 0.77** | 0.86** | 0.80** | 0.89*** | 0.44ns | 0.05ns | 0.05ns |
| **26** | 0.11ns | -0.93ns | 0.17ns | 0.64ns | 0.71ns | 0.86*** | 0.84*** | 0.94*** | 0.84*** |
| **27** | NA | 0.80ns | 0.89** | 0.24ns | -0.02ns | 0.65ns | 0.60ns | 0.47ns | 0.70ns |
| **28** | 0.95ns | 0.88* | 0.60ns | 0.42ns | 0.50ns | 0.60ns | 0.61ns | 0.42ns | 0.40ns |
| **29** | -0.14ns | -0.48ns | -0.72ns | -0.51ns | -0.31ns | 0.17ns | 0.68ns | 0.63ns | 0.30ns |
| **30** | 0.95** | 0.25ns | 0.03ns | 0.19ns | 0.41ns | 0.49ns | 0.55ns | 0.63ns | 0.63ns |
| **31** | NA | NA | NA | 0.74ns | 0.06ns | 0.40ns | -0.03ns | 0.62ns | 0.30ns |
| **32** | NA | NA | -0.57ns | 0.05ns | 0.28ns | -0.05ns | -0.46ns | -0.07ns | -0.10ns |

*^a^* Pearson’s coefficients of correlation r and statistical significance (H_0_: r = 0).

*, **, *** : p-value < 5%, < 1% and < 0.1% ; ns : not significant, *i.e*. p-value > 5%.
