## Supplementary material for "The quasi-universality of nestedness in the structure of quantitative plant-parasite interactions": Suppl Methods 1 and 2; Suppl. Tables 19 to 22: Table S22.docx

**Supplementary Table S22**. Analysis of modularity of plant-parasite interaction matrices with three algorithms, *edge betweenness*, *fast greedy* and *louvain*. Only the three null models S, C2 and R2 that provided the lowest rates of false positive modularity in our performance study (Supplementary Methods 2) are presented.

| **Matrix number** | ***Edge betweenness*** | | | | ***Fast greedy*** | | | | ***Louvain*** | | | |
| --- | --- | --- | --- | --- | --- | --- | --- | --- | --- | --- | --- | --- |
|  | **Modularity score***^a^* | **Null model***^b^* | | | **Modularity score***^a^* | **Null model***^b^* | | | **Modularity score***^a^* | **Null model***^b^* | | |
|  |  | **S** | **C2** | **R2** |  | **S** | **C2** | **R2** |  | **S** | **C2** | **R2** |
| 1 | 0.012 | **1***^c^* | **1** | **0.99** | 0.074 | **1** | **0.99** | 0.51 | 0.073 | **1** | **1** | 1 |
| 2 | 0.022 | **1** | **1** | **0.97** | 0.091 | **1** | **1** | 0.98 | 0.091 | **1** | **1** | 1 |
| 3 | 0.099 | 0.88 | 0.53 | 0.06 | 0.126 | **1** | **0.96** | 0.38 | 0.127 | **1** | 1 | 0.64 |
| 4 | 0 | **1** | **1** | **1** | 0.113 | **1** | **1** | **1** | 0.115 | **1** | **1** | **1** |
| 5 | 0.018 | **1** | **0.01** | **1** | 0.065 | **0.99** | **0** | **0.99** | 0.069 | **1** | **0** | **1** |
| 6 | 0.085 | **1** | **0.99** | **1** | 0.209 | 0.88 | **0.03** | 0.17 | 0.209 | 0.96 | 0.10 | 0.42 |
| 7 | 0.138 | 0.96 | 0.45 | 0.79 | 0.240 | **1** | 0.84 | **1** | 0.240 | **1** | 0.98 | **1** |
| 8 | 0.003 | **1** | **1** | **1** | 0.121 | **1** | **1** | **0.98** | 0.125 | **1** | **1** | 1 |
| 9 | 0 | 1 | 1 | 1 | 0.074 | **1** | **1** | **1** | 0.069 | **1** | **1** | **1** |
| 10 | 0 | **1** | **1** | **1** | 0.073 | **1** | **1** | **1** | 0.073 | **1** | 0.98 | **1** |
| 11 | 0 | **1** | **1** | **1** | 0.085 | **1** | **0.98** | **1** | 0.091 | **1** | 0.93 | **1** |
| 12 | 0.004 | **1** | **1** | 0.50 | 0.041 | **1** | **1** | 0.16 | 0.044 | **1** | **1** | 0.98 |
| 13 | 0.004 | **1** | **1** | **0.95** | 0.061 | **1** | **1** | 0.56 | 0.063 | **1** | **1** | 0.88 |
| 14 | 0 | 1 | 1 | 1 | 0.044 | 0.87 | **0.04** | 0.35 | 0.046 | 0.48 | **0.04** | 0.15 |
| 15 | 0 | **1** | **1** | **1** | 0.089 | **1** | **0.98** | **1** | 0.087 | **1** | **1** | **1** |
| 16 | 0.007 | **1** | 1 | **1** | 0.032 | **1** | **1** | **1** | 0.036 | **1** | **1** | **1** |
| 17 | 0.071 | **1** | **0.99** | **1** | 0.172 | **1** | **1** | **1** | 0.177 | **1** | **1** | **1** |
| 17b*^d^* | 0.040 | **1** | **1** | **0.99** | 0.155 | **1** | **1** | **1** | 0.158 | **1** | **1** | **1** |
| 18 | 0.079 | 1 | 1 | 1 | 0.186 | **1** | **1** | 0.35 | 0.186 | **0.99** | **1** | 0.16 |
| 19 | 0 | **1** | 1 | **1** | 0.028 | **1** | **1** | **1** | 0.031 | **1** | 1 | **1** |
| 20 | 0 | **1** | 1 | 1 | 0.029 | **1** | **1** | **1** | 0.029 | **1** | **1** | **1** |
| 21 | 0.036 | **1** | 0.23 | **1** | 0.110 | **1** | **0** | **1** | 0.109 | **1** | **0.01** | **1** |
| 22 | 0.020 | **1** | **0.98** | **1** | 0.165 | **1** | 0.89 | **1** | 0.165 | **1** | 0.93 | **1** |
| 23 | 0.011 | **1** | **1** | 0.88 | 0.068 | 0.91 | 0.81 | 0.22 | 0.070 | 0.99 | 0.99 | 0.70 |
| 24 | 0.035 | 0.99 | 0.86 | 0.91 | 0.092 | 0.79 | 0.75 | 0.20 | 0.095 | 0.94 | 0.93 | 0.60 |
| 25 | 0.066 | **1** | 0.35 | **1** | 0.167 | **1** | 0.49 | **1** | 0.168 | **1** | 0.84 | **1** |
| 26 | 0 | **1** | **1** | **1** | 0.041 | **1** | **1** | **1** | 0.046 | **1** | 1 | **1** |
| 27 | 0.034 | 0.96 | 0.86 | 0.69 | 0.102 | **1** | **1** | **1** | 0.103 | **1** | **1** | **1** |
| 28 | 0 | **1** | **1** | **1** | 0.151 | **1** | **1** | **1** | 0.153 | **1** | **1** | **1** |
| 29 | 0.008 | **1** | **1** | **1** | 0.069 | **1** | **1** | **1** | 0.069 | **1** | **1** | **1** |
| 30 | 0.043 | **0.98** | 0.91 | 0.91 | 0.087 | **1** | **1** | **1** | 0.092 | **1** | **1** | **1** |
| 31 | 0.013 | **0.98** | **0.97** | 0.65 | 0.062 | **1** | **1** | 0.89 | 0.062 | **1** | **1** | 1 |
| 32 | 0.058 | 0.70 | 0.84 | 0.72 | 0.062 | **0.95** | **0.99** | 0.59 | 0.062 | **1** | **1** | 1 |

*^a^* Maximum of 100 estimates.

*^b^* See Supplementary Methods 2 for details of the null models.

*^c^* Modularity significance: the probability value (p-value) indicates the frequency of null-model matrices showing a strictly higher modularity score than that of the actual matrix. P-values ≤ 0.05 (significant modularity) are in bold on grey cells. Significant anti-modularity, when ≤ 5% of null-model matrices show a strictly lower modularity degree than that of the actual matrix, are indicated in white on black cells. Note that some of the indicated p-values are ≥ 0.95 but do not correspond to significant anti-modularity because the modularity degrees of the actual matrix and of some null-model matrices are identical.

*^d^* Matrix 17b is identical to matrix 17 except that columns entirely made of zero-valued cells and redundant columns were removed.
