## Supplementary material for "The quasi-universality of nestedness in the structure of quantitative plant-parasite interactions": Suppl Methods 1 and 2; Suppl. Tables 19 to 22: TableS20.docx

**Supplementary Table S20.** Parts of phenotypic variance explained by the plant genotype, the parasite strain and their interaction for the analysed datasets.

| **Matrix number** | ω²_plant_*^a^* | ω²_parasite_ | ω²_plant × parasite_ | ω²_plant × parasite_ / ω²_total_ |
| --- | --- | --- | --- | --- |
| **1** | 0.11 | 0.16 | 0.02 NS*^b^* | 0.08 NS |
| **2** | 0.04 | 0.41 | 0 NS | 0 NS |
| **3** | 0.11 | 0.17 | 0.04 | 0.11 |
| **4** | 0.11 | 0.25 | 0.06 | 0.14 |
| **5** | -*^c^* | - | - | - |
| **6** | 0.23 | 0.21 | 0.18 | 0.29 |
| **7** | - | - | - | - |
| **8** | 0.11 | 0.41 | 0.13 | 0.20 |
| **9** | - | - | NS after Soltis et al. (2019) | NS after Soltis et al. (2019) |
| **10** | - | - | - | - |
| **11** | - | - | - | - |
| **12** | 0.07 | 0.49 | 0 NS | 0 NS |
| **13** | 0.12 | 0.41 | 0 NS | 0 NS |
| **14** | 0.33 | 0.14 | 0.18 | 0.28 |
| **15** | 0.28 | 0.12 | 0.09 | 0.18 |
| **16** | - | - | - | - |
| **17** | 0.44 | 0.24 | 0.16 | 0.19 |
| **18** | 0.51 | 0.28 | 0.08 | 0.10 |
| **19** | 0.76 | 0.06 | 0.06 | 0.07 |
| **20** | 0.22 | 0.18 | 0.04 | 0.09 |
| **21 (non nested)** | 0.55 | 0.03 | 0.13 | 0.19 |
| **22** | 0.57 | 0.08 | 0.13 | 0.16 |
| **23** | 0.24 | 0.15 | 0.19 | 0.32 |
| **24** | 0.15 | 0.17 | 0.22 | 0.41 |
| **25** | 0.29 | 0.17 | 0.28 | 0.38 |
| **26** | 0.48 | 0.16 | 0.10 | 0.13 |
| **27** | 0.23 | 0.32 | 0.11 | 0.17 |
| **28** | 0.16 | 0.26 | 0.12 | 0.23 |
| **29** | 0.38 | 0.30 | 0.19 | 0.22 |
| **30** | 0.30 | 0.39 | 0.16 | 0.19 |
| **31** | 0.11 | 0.21 | 0.15 | 0.31 |
| **32 (non nested)** | 0.04 | 0.21 | 0.27 | 0.52 |

*^a^* Unbiased estimator of the part of variance explained by a given effect in the linear model

*pathogenicity ~ ‘parasite strain’ + ‘plant accession’ + ‘parasite strain × plant accession’* calculated as:

ω²_effect_= (SS_effect_ – df_effect_ × MS_error_) / (SS_total_ + MS_error_), where SS : sum of squares, df : number of degrees of freedom and MS : mean squares (Kirk, 1982).

*^b^* NS : the plant × parasite interaction was not significant.

*^c^* - : data not available.
